# Carbendazim Disrupts Cellular Homeostasis and Proteostasis in *Caenorhabditis elegans*: A Global Proteomic Approach

**DOI:** 10.64898/2026.10.12.758533

**Authors:** Aline Castro Silva, Maria Luiza de Lima Vitorino, Emilly de Souza Cordeiro, Helena Oliveira Barcelos, Daniel Balbé Nunes, João Gabriel Barbosa de Luna, Josivan Barbosa de Farias, Roberto Afonso da Silva, José Luiz de Lima Filho, Priscila Gubert, Leonardo Fernandes Fraceto, Simone Pinton, Daiana Silva Ávila

## Abstract

The extensive use of the broad-spectrum fungicide carbendazim (CBZ) has raised increasing environmental concerns due to its persistence and toxicity, leading to regulatory restrictions in several countries. Despite its widespread occurrence, the molecular mechanisms underlying CBZ toxicity in non-target organisms remain poorly understood. Here, we employed quantitative proteomics to characterize the systemic molecular response of *Caenorhabditis elegans* following exposure to a sublethal concentration of CBZ (0.01 mg·mL^-1^) from the L1 larval stage to day 1 of adulthood. A total of 273 differentially abundant proteins were identified in the N2 strain. Functional enrichment analysis revealed coordinated alterations in biological processes associated with protein synthesis, proteostasis, mitochondrial function, energy and lipid metabolism, intracellular trafficking, chromatin organization, and redox homeostasis, indicating extensive cellular reprogramming in response to CBZ exposure. Protein–protein interaction network analysis further identified ribosomal proteins and translation-associated factors as the principal hub proteins, suggesting that disruption of the translational machinery represents a central molecular event underlying CBZ toxicity. Collectively, these findings show that CBZ’s biological effects extend beyond its established role as a microtubule-disrupting fungicide, affecting multiple interconnected cellular pathways involved in maintaining homeostasis. This study provides the first comprehensive proteomic characterization of CBZ toxicity in *C. elegans*, offering new mechanistic insights into its mode of action and highlighting the utility of integrative proteomics for assessing the impact of environmental contaminants on non-target organisms.

Graphical abstract.

*Caenorhabditis elegans* (N2) exposed to carbendazim (0.01 mg·mL^-1^) was subjected to UPLC–mass spectrometry-based proteomic analysis. Carbendazim exposure induced broad proteomic alterations associated with lipid metabolism, protein folding, mitochondrial function, central metabolism and protein synthesis.

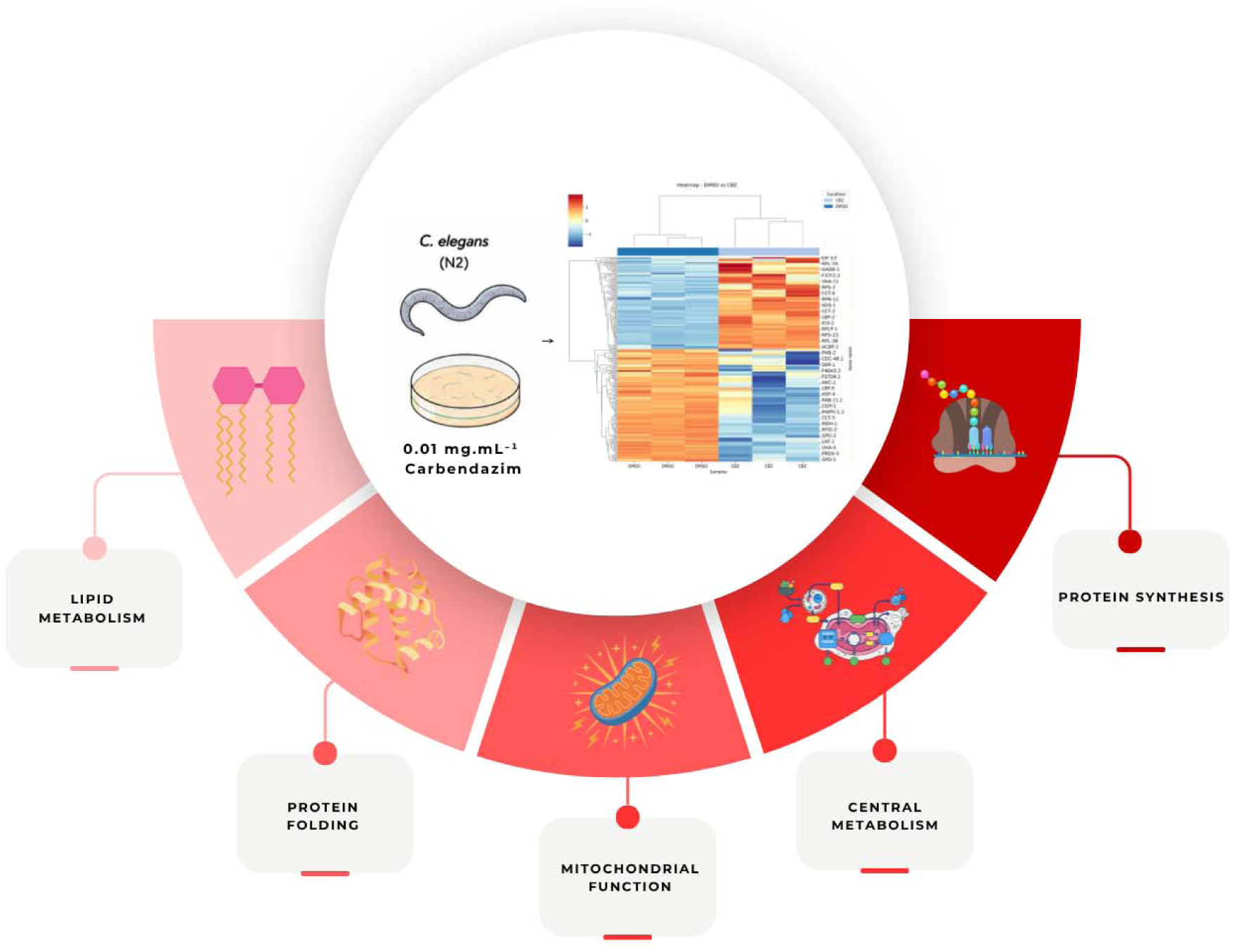

## 1. Introduction

The increase in agricultural production to meet the growing global demand for food, along with the expansion of grain exports and their by-products, has led to the intensive use of pesticides aimed at reducing losses caused by biotic and abiotic factors (Hemathilake & Gunathilake, 2022). Currently, global pesticide consumption reaches approximately 2.6 million tons per year, of which nearly 17% corresponds to fungicides used to control fungal infestations. In parallel, the global pesticide market was valued at approximately USD 84.5 billion in 2019, exhibiting an annual growth rate exceeding 4% since 2015, which highlights the increasing dependence on these chemical inputs in modern agriculture (Aktar et al., 2009; Heinrich Böll Foundation, 2022). Although pesticides play a crucial role in pest and disease control, their excessive and often poorly regulated application has raised serious concerns regarding environmental contamination and risks to human health, as observed for compounds such as carbendazim (CBZ) (Figure S1) (Heinrich Böll Foundation, 2022).

CBZ is a systemic, broad-spectrum benzimidazole-carbamate fungicide that inhibits microtubule synthesis in fungal cells, thereby blocking cell division and fungal proliferation (Bai et al., 2024).It has been widely applied in agriculture on fruits, vegetables, cereals, and flowers. Due to its environmental persistence and associated toxicological risks, CBZ has been banned in countries such as Brazil, the United States, and those in the European Union; however, its use remains permitted in several developing nations, including China, India, and some Latin American countries (De et al., 2014; Y. Wang et al., 2014). In Brazil, CBZ was also banned by the Health Regulatory Agency (ANVISA) in August 2022, with its importation, production, commercialization, and use currently being phased out (ANVISA, 2022a; IDESF, 2022). This decision followed toxicological evaluations indicating potential carcinogenic, mutagenic, and reproductive effects, as well as environmental concerns related to soil and water contamination and impacts on non-target organisms (ANVISA, 2022b). Despite these risks, CBZ and its derivatives have been investigated as potential chemotherapeutic agents due to their ability to inhibit the proliferation of human breast, lung, and colon cancer cells (Lee et al., 2023). However, regardless of its potential biomedical applications, CBZ remains an environmental contaminant capable of affecting non-target organisms. Although its adverse effects on wildlife and ecosystem health have been widely reported, the molecular pathways underlying CBZ-induced toxicity remain poorly understood. In particular, comprehensive proteomic studies investigating the global cellular responses to CBZ exposure in environmentally relevant models are still scarce.

Alternative *in vivo* models such as *Caenorhabditis elegans* have proven highly suitable for mechanistic toxicological investigations, particularly in omics-based approaches. This nematode combines a short life cycle, low maintenance cost, ease of handling, and high genomic homology with humans, while also possessing a well-annotated proteome and conserved molecular pathways (Bezerra et al., 2025; Hunt, 2017; Li & Zhou, 2026; Silva et al., 2025; Xu et al., 2026). These characteristics make *C. elegans* an excellent platform for systems-level toxicological studies, particularly quantitative proteomics, enabling the identification of global protein expression changes, disrupted biological pathways, and molecular networks associated with xenobiotic exposure (Long et al., 2023). Additionally, as a soil-dwelling organism, it represents a relevant model for ecotoxicological studies investigating the impact of pesticides on non-target organisms.

Therefore, investigating the molecular effects of CBZ in non-target organisms is essential to improve our understanding of its mechanisms of toxicity and potential environmental impacts. We hypothesized that sublethal exposure to CBZ induces proteomic alterations in *C. elegans*, which may be associated with locomotor impairment and neurotoxicity. Accordingly, we performed a comprehensive quantitative proteomic analysis to characterize the global protein expression changes induced by CBZ exposure. By integrating proteomic profiling with functional and network analyses, this study provides mechanistic insights into CBZ toxicity that may guide future investigations in more complex biological systems and contribute to the development of safer strategies for its use.

## 2. Material and methods

### 2.1 C. elegans strains and maintenance

Worms were cultivated in Petri dishes containing nematode growth medium (NGM) (Stiernagle, 2006) and fed with *Escherichia coli* OP50 under controlled temperature (20 °C) and humidity (>95%) (Panasonic Healthcare Company of North America, MIR-254-PA). The population was synchronized to obtain all worms at the same larval stage. For this, we used a bleaching solution (10 M NaOH, 1% NaClO, and distilled H_2_O) to break the cuticle of pregnant hermaphrodites and release the eggs. All the strains employed underwent the same process. The strains used were N2 (wild type) and *Escherichia coli* OP50, obtained from the *Caenorhabditis Genetics* Center (CGC, Minnesota, USA).

### 2.2 Exposure protocol

We used 20,000 *C. elegans* synchronized at the L1 stage, which were transferred to Petri dishes (150 mm) containing 80 mL of NGM previously seeded with *E. coli* OP50 and CBZ, for 72 h (L1-day 1 adults). For the analyses, the following groups were used: DMSO control (exposed to 0.6% dimethyl sulfoxide) and CBZ at 0.01 mg.mL^-1^. The concentration of CBZ was selected as a sublethal dose to induce biological responses while preserving animal viability for proteomic analysis (Figure S1). CBZ was obtained from Sigma-Aldrich (St. Louis, MO, USA). All other reagents were of analytical grade and were obtained from local suppliers.

### 2.3 Proteomics analysis

#### 2.3.1 Protein sample preparation

Approximately 20,000 worms of each group were subjected to protein extraction. The extraction was performed by manual maceration using the Sample Grinding KitTM (Cytiva, Marlborough, MA) and using a Lysis buffer (7 M urea, 2 M thiourea, 4% 3-[(3-cholamidopropyl)dimethylammonio]-1-propanesulfonate, 40 mM dithiothreitol (DTT), 0.5% Immobilized pH Gradient Buffer). During extraction, a protease inhibitor cocktail from Promega (Madison, Wisconsin, USA) was added at a protein ratio of 1:50 w/w to the mixture.

The protein concentration was determined using the 2-D Quant KitTM (Cytiva, Marlborough, MA). Afterward, 100 μg of protein from each group was submitted to a 2-D Clean-Up TM Kit (Cytiva, Marlborough, MA) protocol to remove any interfering components, following the manufacturer’s instructions. 100 μg of protein from each group was resuspended in 50 μL of 8 M urea. The proteins were reduced by adding 100 mM DTT at 30 °C for 30 minutes and alkylated with 300 mM Iodoacetamide. Afterward, 50 mM NH_4_HCO_3_ was added to the solution. For protein digestion, sequencing-grade modified trypsin (Madison, Wisconsin, USA) was added at a protein/protein ratio of 1:50 w/w, and the samples were incubated at 37 °C for 18 hours. The tryptic peptides were concentrated at 30 °C on a Speed Vacuum concentrator 5301 Eppendorf (Hamburg, Germany) (Farias et al., 2024).

#### 2.3.2 Liquid chromatography–tandem mass spectrometry (LC-MS/MS) analysis

The samples of tryptic peptides were recovered in 0.1% formic acid at a final concentration of 1 μg μL^-1^. The tryptic peptides were separated using an M-Class ACQUITY UPLC system (Waters Corporation, Milford, MA). This system was equipped with a Single Pump Trap with column: nanoease M/Z symmetry C18 100 A, 5 μm, 180 μm x 20 mm trap column, operating at 0.3 μL min-1, coupled to an analytical nanoease M/Z HSS C18 T3, 100 A, 1.8 μm, 75 μm x 250 mm column operating at 0.3 μL min^-1^ and a temperature of 40 °C. The mobile phase A consisted of water containing 0.1% (v/v) formic acid, while the mobile phase B consisted of acetonitrile containing 0.1% (v/v) formic acid. Peptide elution was performed for 133 min and loading time of 10 min using the following gradient: (I) 0 – 8 min, 95% of A and 5% of B; (II) 8 – 98 min, 60% of A and 40% of B; (III) 98 – 103 min, 15% of A and 85% of B; (IV) 103 – 108 min, 15% of A and 85% of B; (V) 108 – 109 min, 95% of A and 5% of B; (VI) 109 – 133 min, 95% of A and 5% of B (starting conditions).

The system was coupled to a Synapt XS quadrupole Q-ToF mass spectrometer with T-Wave Ion Mobility (Waters Corporation, Wilmslow, UK) operating at a mass resolution of 30,000, Full Width at Half Maximum. The Electrospray Low Flow probe capillary voltage was 3 kV, sampling cone 40 V; source offset 30 V, source temperature 100 °C, and cone gas 50 L min-1. The time-of-flight mass analyzer was externally calibrated with a NaCsI mixture from m/z 50 to 2000. A lock mass reference signal of GluFibrinopeptide B (m/z 785.8426) was sampled every 30 s. Data were acquired in triplicate using the Ultra-Definition Mass Spectrometry in positive ion mode.

#### 2.3.3 Data processing, protein identification, and quantification

Protein identification was performed on LC-MS/MS data using Progenesis QI v 4.7 software (Nonlinear Dynamics) with a revised protein database that includes the *Caenorhabditis elegans* proteome (UniProt_download on 21/11/2023) (Bateman et al., 2023). The parameters for spectra processing and database search were loss of trypsin cleavage, max protein mass 750 KDa, fixed modification of carbamidomethyl to Cysteine (Cys), and variable modification of oxidation to methionine (Met). The search thresholds used were (2) minimum fragments ions matches per peptide, (5) minimum fragments ions match per protein, (2) minimum fragment ion matches per peptide per protein, and (1) false discovery rate (FDR) for protein identification. Relative quantification was performed using the three most abundant peptides for all identified proteins (Hi3 label-free method). Only proteins with frequency scores and confidence intervals greater than 99% were considered acceptable in the database searches. A simplified schematic illustration of the exposure protocol, the extraction process, and the preparation of peptides for digestion, identification, and analysis is shown in Figure 1.

**Figure 1.**
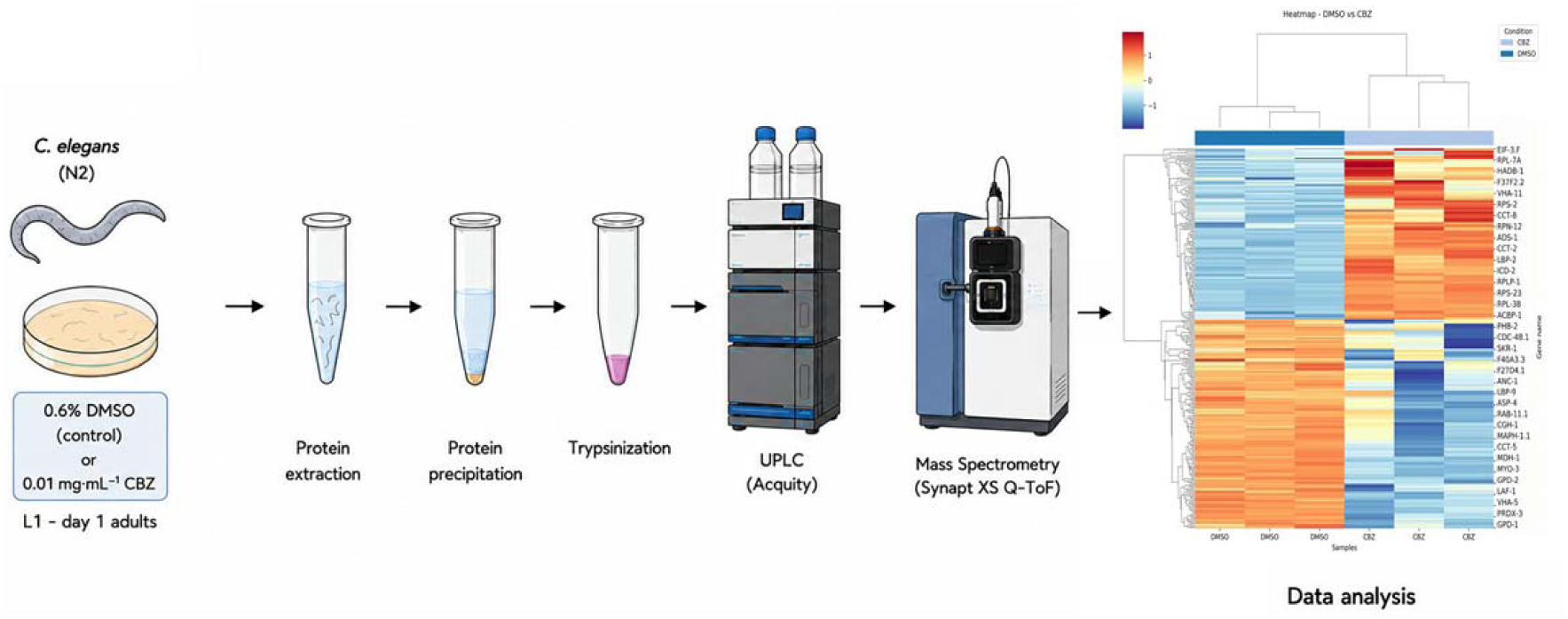
Simplified schematic illustration of the protocol. Exposure, extraction process, and peptide preparation for digestion, identification, and analysis (created by the authors using ChatGPT (OpenAI, 2026), adapted from Bezerra et al., 2025.

#### 2.3.4 Bioinformatics analysis

The Progenesis QI v4.7 software was used to identify, normalize, and extract protein groups. Normalized protein abundance values were employed for all subsequent statistical and bioinformatics analyses. Data processing, visualization, and differential abundance analysis were conducted using the OmicScope pipeline (v1.4.2; Python 3.11) together with the ProteoSpectra script (Bezerra et al., 2025; Reis-de-Oliveira et al., 2024). Statistical evaluation of the proteins was performed using Analysis of Variance (ANOVA), followed by Benjamini-Hochberg multiple-testing correction to control the False Discovery Rate (FDR). Proteins with an adjusted p-value (q-value) < 0.05 and an absolute fold chag (|log FC| ≥ 0.263) were classified as differentially abundant proteins (DAPs) (Zhang et al., 2020).

Based on the Gene Ontology (GO) database, biological process (BP), cellular component (CC), molecular function (MF), anatomy association, KEGG, and Reactome pathway enrichment analyses were performed for the selected DAPs using the DAVID bioinformatics tool (Kanehisa, 2000; Sherman et al., 2024). Statistical significance was established at p-value < 0.05 and FDR < 0.05, and the top 10 enriched terms were plotted (Bezerra et al., 2025; Blaha et al., 2026; Boujenoui et al., 2024; Subhadra et al., 2024). Data visualization was performed using in-house scripts developed in Python 3.8, which utilized the pandas, seaborn, and matplotlib libraries. Code is available at https://github.com/joaoluna-dev/proteomics_c_elegans.

Protein-protein interaction (PPI) networks were mapped using the STRING database (v11.5) based on the identified DAPs. Only interactions meeting a high-confidence score threshold (≥ 0.70) were retained. The resulting interaction networks were imported into Cytoscape (v3.10.1) and customized utilizing the Omics Visualizer plugin (Legeay et al., 2020). To detect highly interconnected functional modules within the networks, cluster analysis was executed via the MCODE plugin using standard criteria (degree cutoff = 2, node score cutoff = 0.2, k-core = 2, and maximum depth = 100). Additionally, topological centrality metrics were evaluated through the CytoHubba plugin to identify hub proteins, filtering the top 10 highest-ranked nodes (Chin et al., 2014).

## 3. Results and Discussion

### 3.1 Proteomic insights into CBZ-induced cellular responses in C. elegans

The intensification of agricultural practices to meet the growing global demand for food has increased reliance on agrochemicals, raising significant environmental and biological concerns (Zhou et al., 2025). CBZ is a widely used systemic fungicide with a well-documented toxicological profile, including endocrine disruption, reproductive toxicity, and neurotoxicity in non-target organisms (Zhou et al., 2023). Using *C. elegans* as a well-established toxicological model with high genetic conservation and suitability for *in vivo* neurobiological analyses (Barbosa et al., 2025; Chatterjee, 2025), we investigated the effects of a sublethal CBZ concentration (0.01 mg·mL^-1^) on nematodes (Figure S2). This concentration is lower than those associated with agricultural applications against white mold (*Sclerotinia sclerotiorum*). Formulations containing 50 g of CBZ per 100 mL, registered in Argentina, are applied at 50–100 cm³·100□L□¹ of water, corresponding to approximately 0.25–0.50 mg.mL^-1^ of CBZ in the spray solution (Nufarm S.A.; Formulagro n.d.). Higher application concentrations were also permitted for certain 500 g·L□¹ CBZ formulations previously registered in Brazil (Nortox S.A.; CCAB Agro S.A., 2009). Our results demonstrate that CBZ exposure induces extensive proteomic remodeling, with proteins involved in translation and protein synthesis emerging as the predominant molecularly affected processes. Collectively, these findings challenge the assumption that sublethal CBZ exposure causes only limited molecular effects in non-target organisms and provide new mechanistic insights into its toxicity.

A volcano plot was generated to visualize the differential protein abundance, applying fold-change (FC) thresholds of ≥1.2 or ≤−1.2 and a significance threshold of p≤0.05 (Figure 2A). A total of 512 proteins were identified across all samples. Proteins that did not present quantified values in all technical replicates were excluded from subsequent analyses. After this filtering step, 496 proteins remained and were used for all statistical and bioinformatics analyses. Differential abundance analysis identified 273 DAPs, comprising 123 upregulated and 150 downregulated proteins in worms exposed to CBZ (0.01 mg.mL^-1^) compared to the DMSO (0.6%) control (Figure 2B) (Table S1). The heatmap revealed a clear separation between the experimental groups, demonstrating a distinct proteomic profile associated with CBZ exposure (Figure 2D). Functional annotation of these proteins indicated enrichment of processes related to protein homeostasis, energy metabolism, vesicular trafficking, cytoskeletal organization, and cellular stress responses.

**Figure 2.**
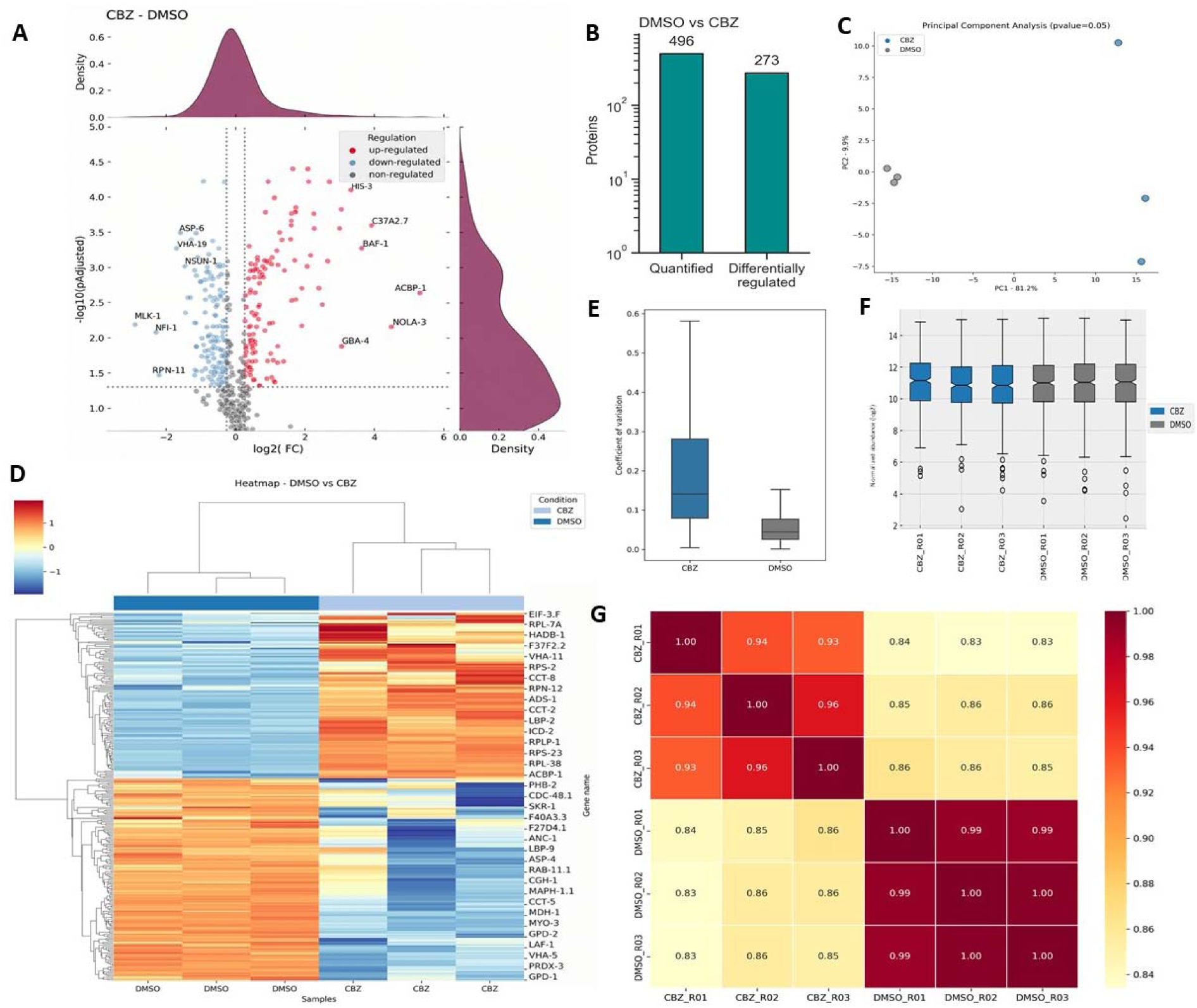
Proteomic analysis of the N2 *C. elegans* strain. (A) Volcano plot of the proteomic profile of N2 exposed to CBZ compared to control (DMSO), with Log2 FC of ≥ 1.2 and ≤ −1.2 and p ≤ 0.05. (B) Data of quantified and differentially regulated proteins. (C) Principal component analysis of data. (D) Heatmap of differentially expressed proteins with repercussions on N2 exposed to CBZ compared to control (DMSO). (E) Coefficient of variation of proteomics replicates. (F) Data normalization. (G) The correlation coefficient between study replicates.

Principal Component Analysis (PCA) revealed a clear separation between the experimental groups (Figure 2C). For the N2 strain, the first principal component (PC1) accounted for 81.2% of the total variance, whereas the second principal component (PC2) explained 9.9%. The distribution of protein coefficients of variation (CVs) is shown in Figure 1C, with outliers removed for visualization purposes. The median CV across biological replicates was 14.1% for the CBZ group and 4.4% for the DMSO control group (Figure 2E). Notably, 62.5% of the proteins in the CBZ group and 96.2% in the DMSO group exhibited CV values below 20%, indicating good reproducibility of the proteomic measurements, consistent with previous proteomic studies in *C. elegans* (Xia et al., 2018). The higher CV values observed for a subset of proteins may be attributed to differences in sampling times between analytical runs and/or the limited detection sensitivity of low-abundance proteins by tandem mass spectrometry (MS/MS). Data normalization was performed to ensure accurate comparisons across samples, resulting in consistent mean protein abundances and reduced error ranges among all conditions and biological replicates (Figure 2F). A heatmap was generated to visually explore expression patterns and variability among the samples (Figure 2D). Pearson correlation analysis was performed to assess pairwise correlations among samples and to identify potential outliers, technical variations, reproducibility issues, and normalization inconsistencies. All experimental groups exhibited strong correlations among biological replicates within the same group, demonstrating the robustness and consistency of the proteomic dataset (Figure 2G).

CBZ exposure induced marked alterations in proteins associated with translation and ribosome biogenesis in *C. elegans*. The upregulation of C37A2.7, encoding the 60S acidic ribosomal protein P2, together with increased abundance of NOLA-3, a nucleolar protein involved in ribosome biogenesis and ribonucleic acid (RNA) processing, suggests enhanced or compensatory regulation of the translational machinery under stress (Chen & Huang, 2001). Consistently, several ribosomal and translation-related proteins, including EIF-3.F, RPL-7A, RPS-2, RPS-23, RPL-38, and RPLP-1, were upregulated. This coordinated response may reflect increased protein synthesis to replace damaged proteins and support cellular adaptation to CBZ-induced stress (Campos et al., 2020; Itani et al., 2021; Ni & Buszczak, 2023). The upregulation of ICD-2, encoding a subunit of the nascent polypeptide-associated complex (NAC), further supports this interpretation, as NAC participates in the proper targeting and early processing of nascent proteins and may contribute to co-translational protein quality control (Gamerdinger, 2016).

In parallel with alterations in protein synthesis, CBZ affected multiple components of the proteostasis network. The increased abundance of RPN-12, a non-ATPase regulatory subunit of the 26S proteasome, suggests remodeling of the proteasomal machinery in response to proteotoxic stress. Previous studies have shown that RPN-12 plays specific roles in *C. elegans* germline development that are independent of overall proteasomal proteolytic activity, with its loss leading to reproductive defects (Fernando et al., 2021). In contrast, RPN-11, another essential proteasomal component, was downregulated, indicating that proteasome-associated responses may be selectively regulated rather than uniformly activated (Hegde et al., 2025; Kandel et al., 2024). The concomitant downregulation of SKR-1, a component of SCF ubiquitin ligase complexes (Schulman et al., 2000), and CDC-48.1, an AAA+ ATPase involved in protein quality control and unfolded protein response (UPR) (Jeon & Ronai, 2026), further supports the involvement of ubiquitin-dependent protein turnover and protein quality control pathways in the cellular response to CBZ. Collectively, these changes suggest a heterogeneous remodeling of the proteostasis network that may compromise the cellular capacity to maintain protein quality under prolonged toxic stress.

The chaperones involved in protein folding, CCT-2 and CCT-8, components of the CCT/TRiC chaperonin complex, were upregulated, whereas CCT-5 was downregulated. Because CCT/TRiC is essential for the adenosine triphosphate (ATP)-dependent folding of cytoskeletal proteins, particularly actin and tubulin (Ge staut et al., 2022), these opposing changes suggest selective remodeling of the chaperone machinery in response to CBZ exposure. This finding is particularly relevant given that CBZ is a microtubule-disrupting fungicide that interacts with β-tubulin (Vela-Corcía et al., 2018). Altered CCT expression may therefore represent a compensatory response to cytoskeletal stress, while reduced abundance of specific subunits could compromise protein-folding capacity and contribute to cellular vulnerability.

Additional alterations in NSUN-1, an RNA methyltransferase involved in ribosomal RNA modification (Heissenberger et al., 2020; Schosserer et al., 2015), and NFI-1, a transcription factor associated with developmental and metabolic regulation (Andrade et al., 2016; Olaseni et al., 2025), indicate that CBZ-induced effects extend beyond protein degradation and folding to broader regulation of RNA metabolism and gene expression. Together, these findings reveal coordinated but heterogeneous remodeling of translation, ribosome biogenesis, protein folding, and degradation pathways, suggesting that CBZ exposure places substantial pressure on cellular proteostasis.

Importantly, disruption of these pathways may have functional consequences. Previous studies in different *in vivo* models have reported behavioral and locomotor alterations following CBZ exposure (Andrade et al., 2016; Olaseni et al., 2025). Consistent with these findings, studies in *C. elegans* have shown alterations in locomotion following exposure to CBZ at concentrations lower than those used in the present study, together with increased reactive oxygen species (ROS) levels (Li et al., 2020). These findings suggest that CBZ toxicity may affect neuromuscular function and cellular redox homeostasis even at relatively low exposure concentrations. Although a direct causal relationship cannot be established from the present proteomic data, the alterations in proteostasis identified here, particularly those involving the proteasome, CCT/TRiC complex, and cytoskeletal protein folding, may provide mechanistic links to the cellular and neuromuscular dysfunction previously associated with CBZ exposure.

Marked alterations in energy and lipid metabolism, together with changes in mitochondrial proteins, were identified in the proteomic profile of *C. elegans* following CBZ exposure. Several proteins involved in central carbon metabolism, including GPD-1, GPD-2, and MDH-1, were downregulated, suggesting disruption of glycolytic, glycerol, and tricarboxylic acid cycle pathways (Lionaki et al., 2022). Consistent with impaired mitochondrial maintenance, PHB-2, a core component of the mitochondrial prohibitin complex involved in mitochondrial biogenesis and integrity (Yoshinaka et al., 2019), and ASP-6, a mitochondrial protease involved in protein turnover and quality control (Yamano et al., 2024), were also reduced. Together, these changes suggest compromised mitochondrial homeostasis and metabolic adaptation following CBZ exposure.

In contrast, the upregulation of HADB-1, a mitochondrial β-oxidation enzyme, and ACBP-1, which encodes an acyl-CoA-binding protein involved in intracellular fatty acid trafficking, may reflect a compensatory shift toward lipid utilization to meet increased energetic demands during stress (Elle et al., 2011; Zhang et al., 2024). Similarly, increased LBP-2 may contribute to fatty acid sequestration and protection against lipid-mediated cellular damage. These responses, together with the altered abundance of LBP-9, F40A3.3, and GBA-4, indicate extensive remodeling of lipid homeostasis, potentially affecting fatty acid trafficking, membrane lipid composition, and sphingolipid metabolism (Clark et al., 2023).

The observed lipid alterations are consistent with previous evidence that CBZ can disrupt lipid metabolism in *C. elegans*, including increased fat accumulation through modulation of the DAF-2/AAK-2 pathway and Δ9-desaturases FAT-6 and FAT-7 (Li & Zhou, 2026). In mammalian models, CBZ exposure has likewise been associated with weight gain, visceral fat accumulation, dyslipidemia, hepatic oxidative stress, and gut microbiota alterations (Bao et al., 2023; Jin et al., 2018; Ma et al., 2023). Collectively, these findings suggest that CBZ induces coordinated remodeling of mitochondrial and lipid metabolism, combining signs of metabolic dysfunction with compensatory mechanisms aimed at maintaining energy homeostasis under toxic stress.

Among the processes altered by CBZ in *C. elegans*, antioxidant defense and stress signaling showed notable changes, consistent with impaired redox homeostasis. Notably, the downregulation of PRDX-3, a mitochondrial peroxiredoxin involved in ROS detoxification, may indicate reduced mitochondrial antioxidant capacity and increased susceptibility to oxidative damage (Sule et al., 2022). This finding is consistent with previous studies associating CBZ toxicity with increased ROS and reactive nitrogen species (RNS) production, lipid peroxidation, and impaired antioxidant defenses in different experimental models (Abolaji et al., 2017; Li et al., 2020; Olaseni et al., 2025b; Rajeswary et al., 2007). Thus, reduced PRDX-3 abundance may contribute to oxidative stress and mitochondrial dysfunction following CBZ exposure.

The downregulation of MLK-1, a MAP3K involved in stress-responsive signaling and environmental adaptation, further suggests alterations in pathways required for an effective cellular response to toxic stress (Brancho et al., 2005). In addition, the upregulation of ADS-1, involved in purine metabolism, may reflect increased nucleotide demand associated with stress-induced cellular remodeling (De Vitto et al., 2021). Together, these alterations indicate that CBZ exposure affects both antioxidant defenses and cellular stress-response pathways.

CBZ exposure also affected proteins involved in chromatin and nuclear organization. The upregulation of HIS-3, encoding histone H3, suggests changes in chromatin dynamics and transcriptional regulation in response to cellular stress (Fabrizio et al., 2019; Filipowicz & Allard, 2025). This finding is consistent with previous evidence that CBZ can disrupt epigenetic regulation, including histone modifications and deoxyribonucleic acid (DNA) methylation, in mammalian models (Liu et al., 2019). Similarly, increased BAF-1, which participates in nuclear envelope organization and chromatin-associated processes, may indicate alterations in nuclear architecture (Montes de Oca et al., 2009) Together, the changes in HIS-3 and BAF-1 suggest coordinated remodeling of chromatin and nuclear organization following CBZ exposure.

Alterations in cytoskeletal and intracellular trafficking proteins further indicate disruption of cellular architecture. The downregulation of ANC-1, a component of the LINC complex linking the cytoskeleton to the nucleus, suggests changes in nuclear–cytoskeletal interactions (Chang et al., 2015), while reduced MAPH-1.1, a microtubule-associated protein, may further compromise microtubule organization and stability, consistent with the established action of CBZ on microtubule dynamics (Yenjerla et al., 2009). In addition, decreased RAB-11.1, a key regulator of endosomal recycling and vesicular trafficking, suggests impaired membrane transport and protein sorting (Winter et al., 2012).

Vacuolar ATPase components were also differentially regulated, with VHA-19 (Colacurcio & Nixon, 2016; Song et al., 2020) and VHA-5 (Eaton et al., 2021) downregulated and VHA-11 (Chung et al., 2019; Song et al., 2020) upregulated. This contrasting pattern suggests remodeling of proton transport and lysosomal function rather than uniform activation or inhibition of these pathways. Given the essential role of V-ATPase in organelle acidification and lysosomal function (Dong et al., 2022), these alterations may influence lysosomal degradation, autophagy, and cellular clearance processes under CBZ-induced stress. Consistent with this possibility, the downregulation of ASP-4, which encodes a lysosomal protease, further suggests a potential reduction in lysosomal proteolytic capacity (Liang et al., 2020). Beyond protein and organelle homeostasis, CBZ exposure also affected RNA metabolism, as indicated by the downregulation of the RNA helicases CGH-1 and LAF-1. These proteins are involved in RNA remodeling, mRNA metabolism, and translational regulation, suggesting that CBZ-induced stress may extend to post-transcriptional mechanisms that contribute to the maintenance of RNA homeostasis and gene expression (Lang et al., 2024).

Overall, the proteomic profile of *C. elegans* exposed to CBZ revealed widespread alterations across multiple interconnected cellular processes, including protein synthesis and proteostasis, mitochondrial function, energy and lipid metabolism, redox homeostasis, intracellular trafficking, cytoskeletal integrity, chromatin organization, and RNA regulation. Together, these changes indicate that CBZ toxicity extends beyond its well-established effects on microtubule dynamics, affecting fundamental mechanisms involved in protein quality control, energy balance, cellular signaling, and maintenance of homeostasis. This integrated response highlights the multifaceted nature of CBZ-induced toxicity and suggests that coordinated disruption of these cellular pathways may contribute to the physiological and functional consequences of CBZ exposure.

### 3.2 Enrichment analysis: GO Enrichment

To elucidate the functional roles of the DAPs, a Gene Ontology (GO) enrichment analysis was performed. Applying the thresholds of p-value < 0.05 and False Discovery Rate (FDR) < 0.05, the results revealed a highly significant and coordinated enrichment in pathways associated with protein synthesis and ribosomal machinery. Within the Biological Process (BP) category, “translation” emerged as the most heavily enriched term (FDR = 7.72e-28), driven by a substantial cluster of ribosomal proteins and eukaryotic translation initiation/elongation factors. Concordantly, the Cellular Component (CC) analysis showed a massive representation of the “ribonucleoprotein complex” (FDR = 1.11e-30), underscoring the predominant subcellular localization of these proteins.

In the Molecular Function (MF) category, the analysis further corroborated these findings, with “structural constituent of ribosome” appearing as the most significant term (FDR = 2.61e-21). Additionally, functions related to protein processing, such as “ATP-dependent protein folding chaperone,” were also significantly enriched. The top 10 most significant results across the GO categories were isolated and visualized, illustrating that the primary molecular impact among the selected DAPs is directed toward the structural and functional regulation of the translation apparatus. Figure 3 shows the top 10 enrichment terms for each GO analysis.

**Figure 3.**
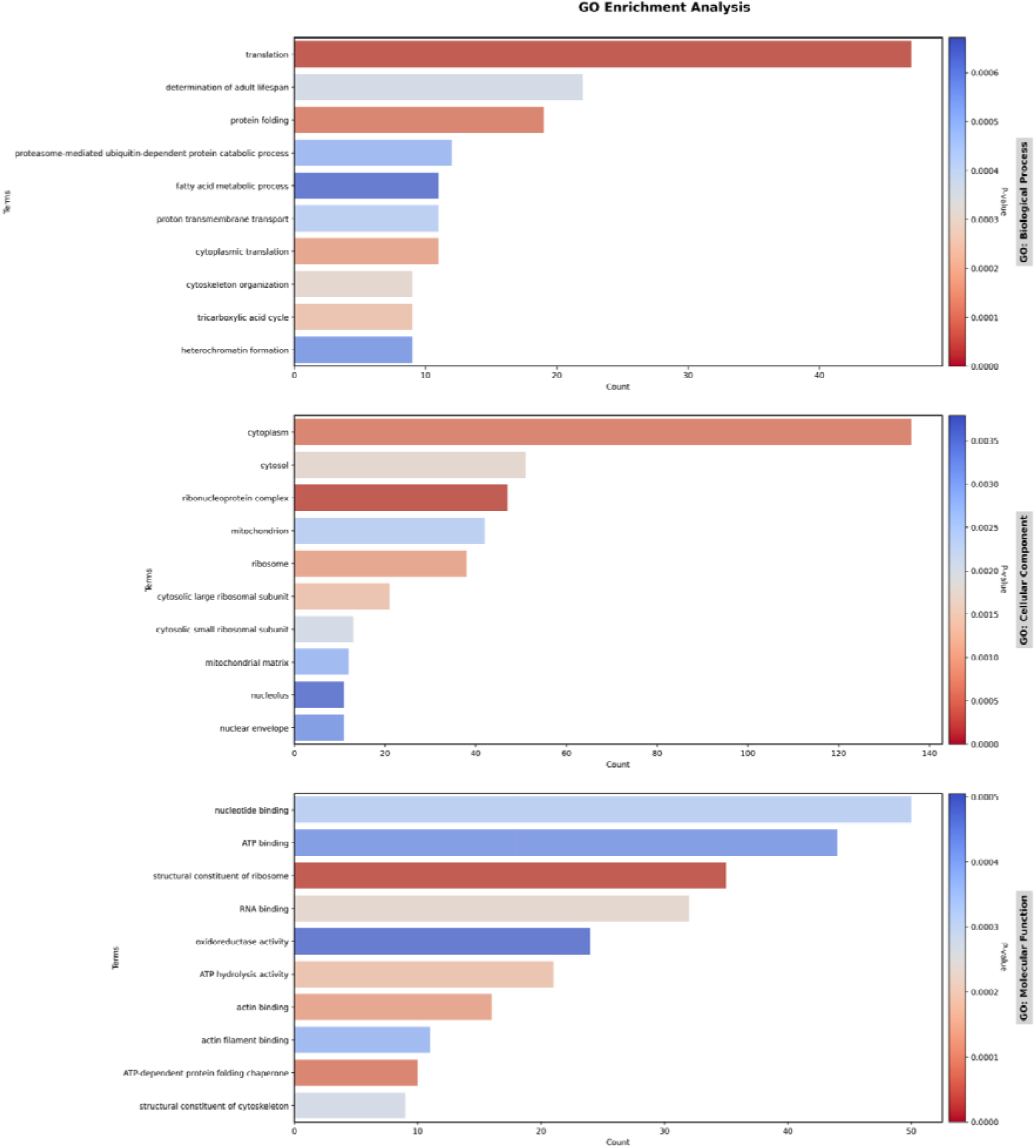
Gene Ontology (GO) enrichment analysis of the differentially expressed proteins. The graph illustrates the top 10 most significantly enriched terms across the Biological Process (BP), Cellular Component (CC), and Molecular Function (MF) categories. Statistical significance was defined by a p-value ≤ 0.05 and a False Discovery Rate (FDR) ≤ 0.05.

### 3.3 Pathway enrichment

To further delineate the metabolic and signaling cascades modulated by the DAPs, pathway enrichment analyses were conducted utilizing the KEGG and Reactome databases. Consistent with the functional ontology findings, networks linked to protein synthesis and homeostatic maintenance were robustly overrepresented. Reactome analysis highlighted “Ribosome-associated quality control” as the most significantly enriched pathway (FDR = 1.36e-27). The KEGG database independently corroborated this biological finding, with “Ribosome” emerging as its top enriched term (FDR = 7.10e-13), underscoring a prominent structural and functional regulation of the translational apparatus.

Beyond the translational machinery, the enriched networks revealed a profound impact on core cellular energetics and metabolism. Under the strict threshold of p-value < 0.05 and FDR < 0.05, KEGG pathway mapping identified substantial enrichment in “Carbon metabolism” (FDR = 3.32e-11), “Citrate cycle (TCA cycle)” (FDR = 2.06e-8), and “Oxidative phosphorylation” (FDR = 3.61e-8). These metabolic findings were strongly supported by the Reactome data, which demonstrated a massive convergence on “The tricarboxylic acid cycle (TCA) and respiratory electron transport” (FDR = 6.48e-20). Additionally, degradative and proteolytic systems, such as the “Proteasome” (KEGG FDR = 6.46e-5), were significantly represented. As visually plotted in the top 10 results, the collective pathway data indicate a comprehensive cellular reprogramming directed toward energy production, carbon metabolism, and protein turnover. Figure 4 illustrates the results of the pathway enrichment analysis.

**Figure 4.**
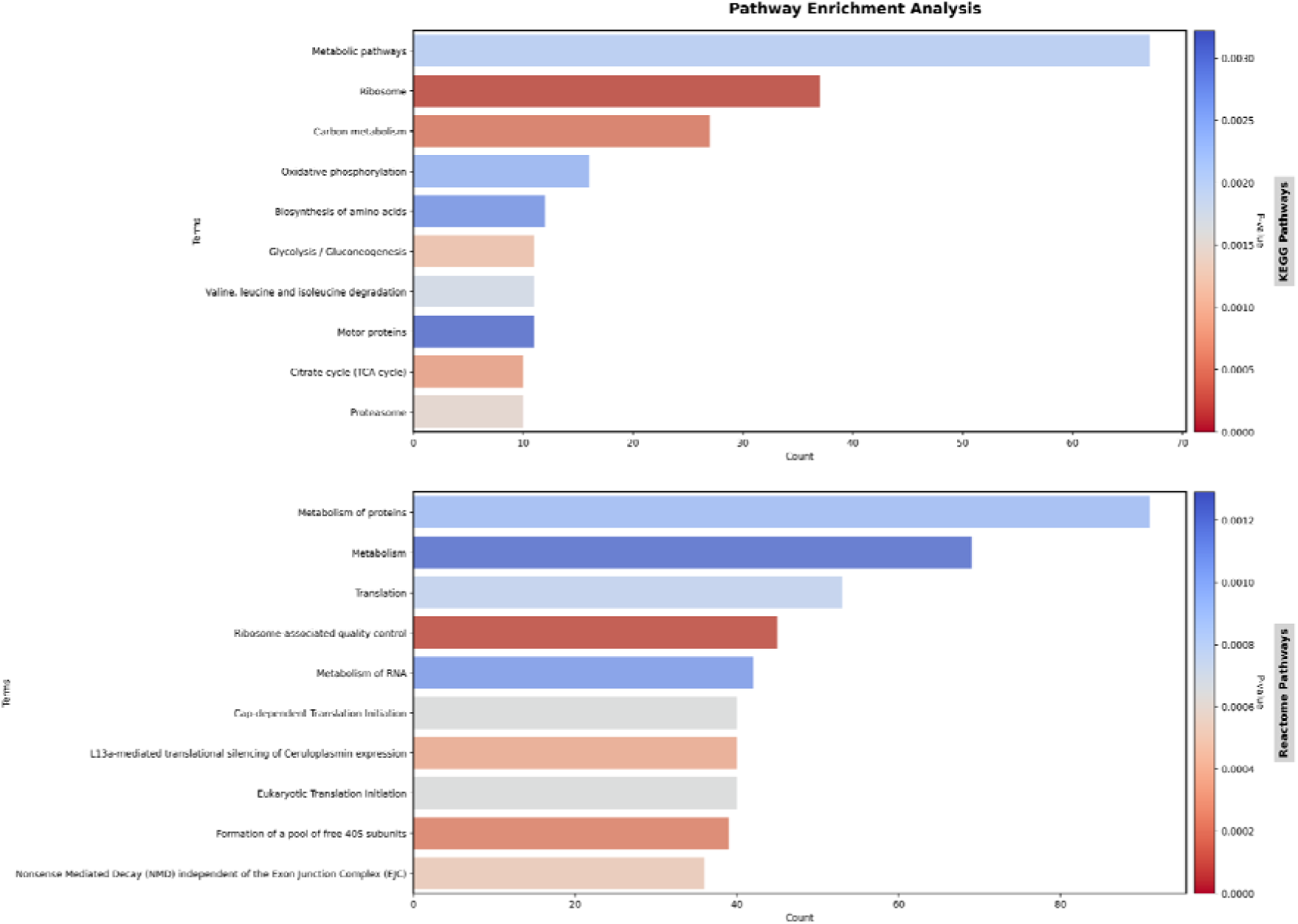
Pathway enrichment analysis of the differentially expressed proteins. The top 10 most significantly enriched metabolic and signaling pathways based on the KEGG and Reactome databases. Statistical thresholds were set at a p-value ≤ 0.05 and a False Discovery Rate (FDR) ≤ 0.05.

### 3.4 PPI Network Analysis Identifies Key Hub Proteins Associated with Protein Synthesis

To identify the key regulatory proteins within the PPI network, hub analysis was performed using three topological algorithms implemented in CytoHubba, including Closeness Centrality (**C_C_**), Degree Centrality, and Maximal Clique Centrality (MCC). Although each method evaluates node importance based on different network properties, all three approaches consistently identified proteins associated with the translational machinery as the major hubs of the network, highlighting protein synthesis as the primary molecular process affected by CBZ exposure (Table 1).

**Table 1.**
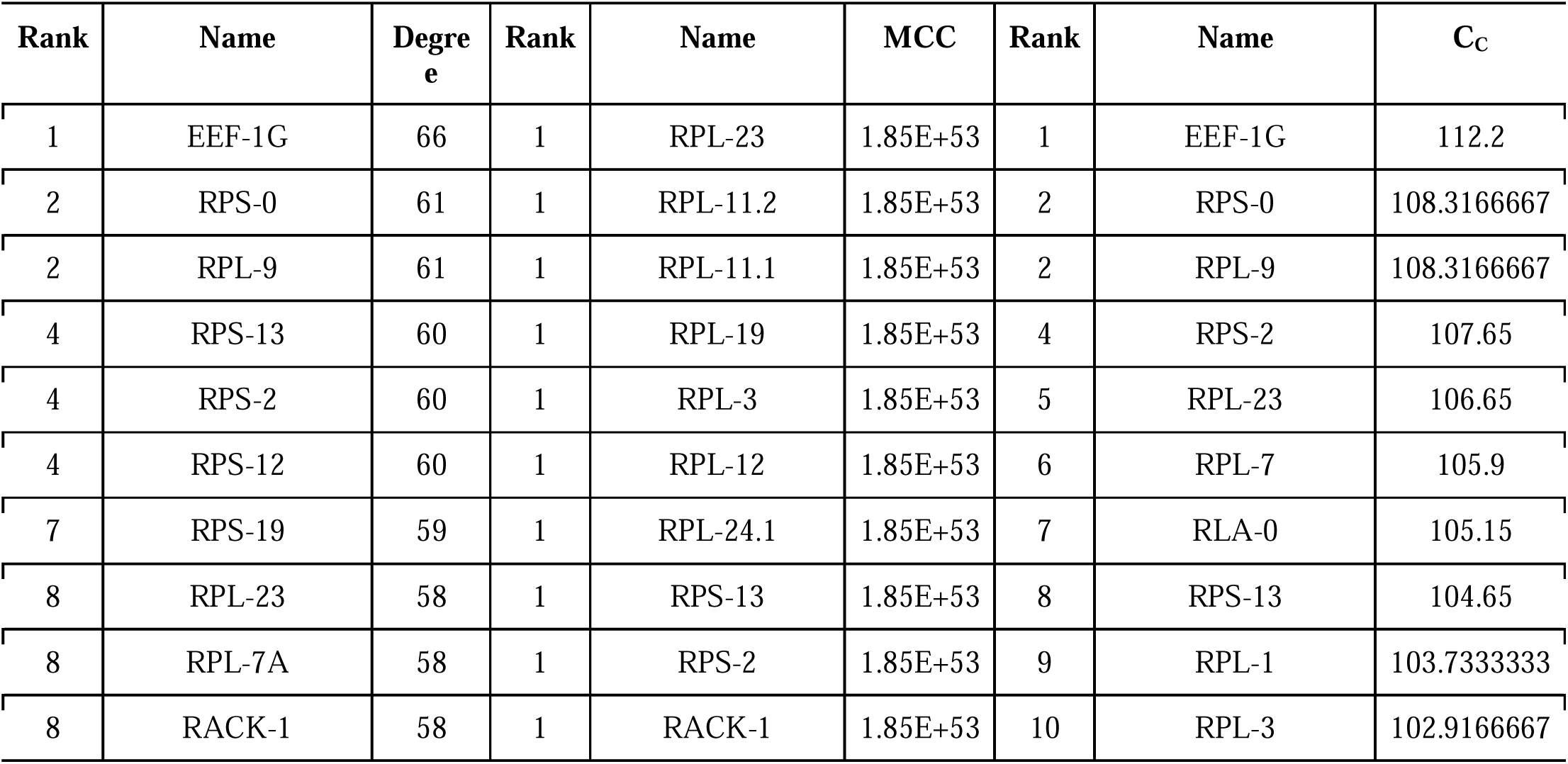
Top 10 proteins with the highest degree (hub genes), MCC, and C_C_ in the *C. elegans* N2.

| Rank | Name | Degree | Rank | Name | MCC | Rank | Name | $C_C$ |
| --- | --- | --- | --- | --- | --- | --- | --- | --- |
| 1 | EEF-1G | 66 | 1 | RPL-23 | 1.85E+53 | 1 | EEF-1G | 112.2 |
| 2 | RPS-0 | 61 | 1 | RPL-11.2 | 1.85E+53 | 2 | RPS-0 | 108.3166667 |
| 2 | RPL-9 | 61 | 1 | RPL-11.1 | 1.85E+53 | 2 | RPL-9 | 108.3166667 |
| 4 | RPS-13 | 60 | 1 | RPL-19 | 1.85E+53 | 4 | RPS-2 | 107.65 |
| 4 | RPS-2 | 60 | 1 | RPL-3 | 1.85E+53 | 5 | RPL-23 | 106.65 |
| 4 | RPS-12 | 60 | 1 | RPL-12 | 1.85E+53 | 6 | RPL-7 | 105.9 |
| 7 | RPS-19 | 59 | 1 | RPL-24.1 | 1.85E+53 | 7 | RLA-0 | 105.15 |
| 8 | RPL-23 | 58 | 1 | RPS-13 | 1.85E+53 | 8 | RPS-13 | 104.65 |
| 8 | RPL-7A | 58 | 1 | RPS-2 | 1.85E+53 | 9 | RPL-1 | 103.7333333 |
| 8 | RACK-1 | 58 | 1 | RACK-1 | 1.85E+53 | 10 | RPL-3 | 102.9166667 |

The Closeness Centrality analysis (Table 1; Figure S2A) identified RLA-0, RPL-7, RPL-1, RPS-13, RPS-0, RPS-2, RPL-23, RPL-3, RPL-9, and EEF-1G as the most central proteins. Closeness centrality measures the average shortest path between one node and all others in the network, indicating proteins that can rapidly influence multiple biological pathways (Hornung & Terrapon, 2022). The predominance of ribosomal proteins, along with the translation elongation factor EEF-1G, suggests that translational regulation occupies a strategic position in coordinating the cellular response to CBZ-induced stress.

Similarly, the Degree Centrality analysis (Table 1; Figure S2B) identified RACK-1, RPL-7A, RPS-19, RPS-12, RPS-13, RPS-2, RPS-0, RPL-23, RPL-9, and EEF-1G as the proteins with the highest number of direct interactions. Proteins exhibiting high degree values function as major interaction hubs and are essential for maintaining network connectivity (Y. Zhang et al., 2013). Among these, RACK-1 is particularly noteworthy because it acts as a scaffold protein associated with the 40S ribosomal subunit, regulating translation initiation and integrating multiple intracellular signaling pathways (Gallo et al., 2018). Its identification as one of the principal hubs suggests that CBZ-induced stress may involve extensive modulation of translational control mechanisms rather than simple changes in ribosome abundance.

The Maximal Clique Centrality (MCC) analysis (Table 1; Figure S2C), which preferentially identifies proteins located within highly interconnected subnetworks and is considered one of the most sensitive algorithms for detecting essential proteins (Wang et al., 2022), revealed RPL-24.1, RPL-11.2, RACK-1, RPS-13, RPL-19, RPS-2, RPL-23, RPL-12, RPL-11.1, and RPL-3 as the top-ranked hubs. The predominance of large ribosomal subunit proteins identified by MCC indicates that these proteins occupy structurally critical positions within the network, where their disruption would likely result in extensive loss of connectivity and functional instability.

Notably, several proteins, including RPS-2, RPS-13, RPL-23, RPL-3, RPL-9, EEF-1G, and especially RACK-1, were identified by more than one algorithm, reinforcing their biological relevance and robustness as central regulators of the CBZ response. The recurrence of these proteins across different topological methods strongly suggests that they represent conserved hubs whose modulation may propagate effects throughout multiple cellular pathways.

From a biological perspective, the predominance of ribosomal proteins among the identified hubs indicates that protein translation represents the principal target of CBZ-induced cellular remodeling. Ribosomes are highly energy-demanding molecular complexes whose regulation is closely linked to stress adaptation. Under toxic conditions, translational reprogramming allows cells to suppress general protein synthesis while selectively translating stress-response proteins involved in detoxification, protein folding, antioxidant defense, and cellular survival (Han et al., 2020).

Furthermore, the identification of EEF-1G and RACK-1 as central nodes suggests that CBZ affects not only the structural composition of ribosomes but also the regulatory mechanisms controlling translation elongation, initiation, and signal integration. RACK-1 is recognized as an evolutionarily conserved scaffold protein that coordinates ribosome-associated signaling, stress responses, and selective mRNA translation (Kolosov et al., 2025a), while EEF-1G participates in the elongation factor complex responsible for accurate protein synthesis (Xu et al., 2022). Alterations in these proteins may therefore have widespread consequences on cellular homeostasis.

In general, the hub analysis demonstrates that the response of *C. elegans* to CBZ is centered on a highly interconnected translational network. The identification of ribosomal proteins and translation regulators as the principal hubs indicates that perturbation of protein synthesis constitutes one of the earliest and most critical events triggered by CBZ exposure. Because hub proteins are essential for maintaining network integrity, their dysregulation is expected to propagate throughout the interactome, affecting multiple downstream biological processes including proteostasis, energy metabolism, stress adaptation, and cell survival.

### 3.5 Protein-protein interaction network

The protein-protein interaction (PPI) network analysis revealed that exposure of *C. elegans* N2 to CBZ (0.01 mg mL^-1^) induced coordinated alterations in multiple biological pathways, indicating a systemic cellular response rather than isolated changes in individual proteins. The resulting PPI network comprised 263 nodes and 2019 edges (Figure 5) and was organized into several highly interconnected clusters, demonstrating that proteins participating in common biological processes are co-regulated following CBZ exposure (Figure 6). MCODE identified 12 modules, of which considered significant with MCODE Score ≥ 5 and generated separate modules from the native network. An analysis of modules in PPI was performed to help us understand the dynamics of *C. elegans* PPI networks. The modules represent groups of proteins that tend to interact with each other more frequently (Guimarães et al., 2018).

**Figure 5.**
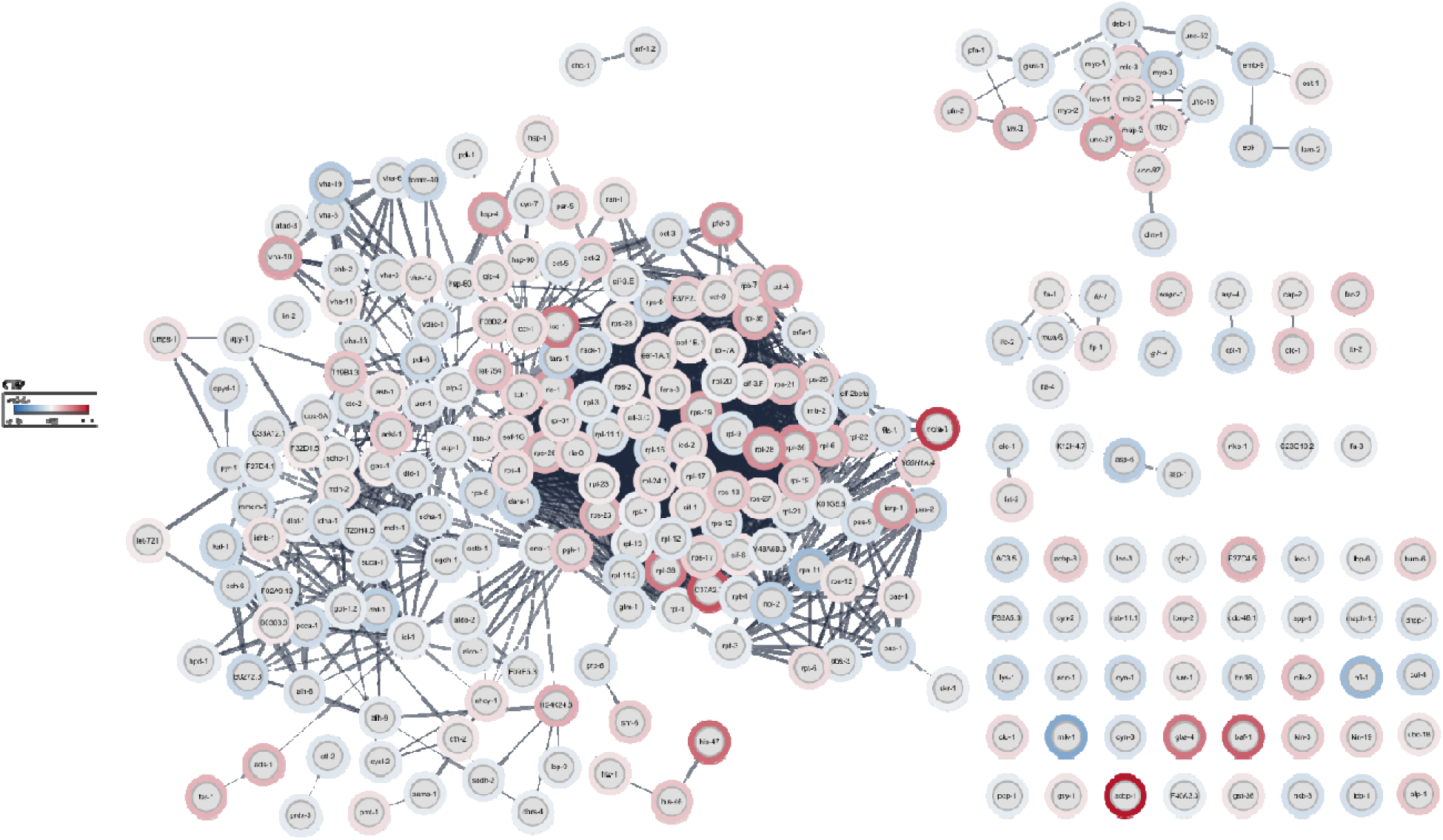
Overall PPI network containing all DAPs from the N2 exposed to CBZ.

**Figure 6.**
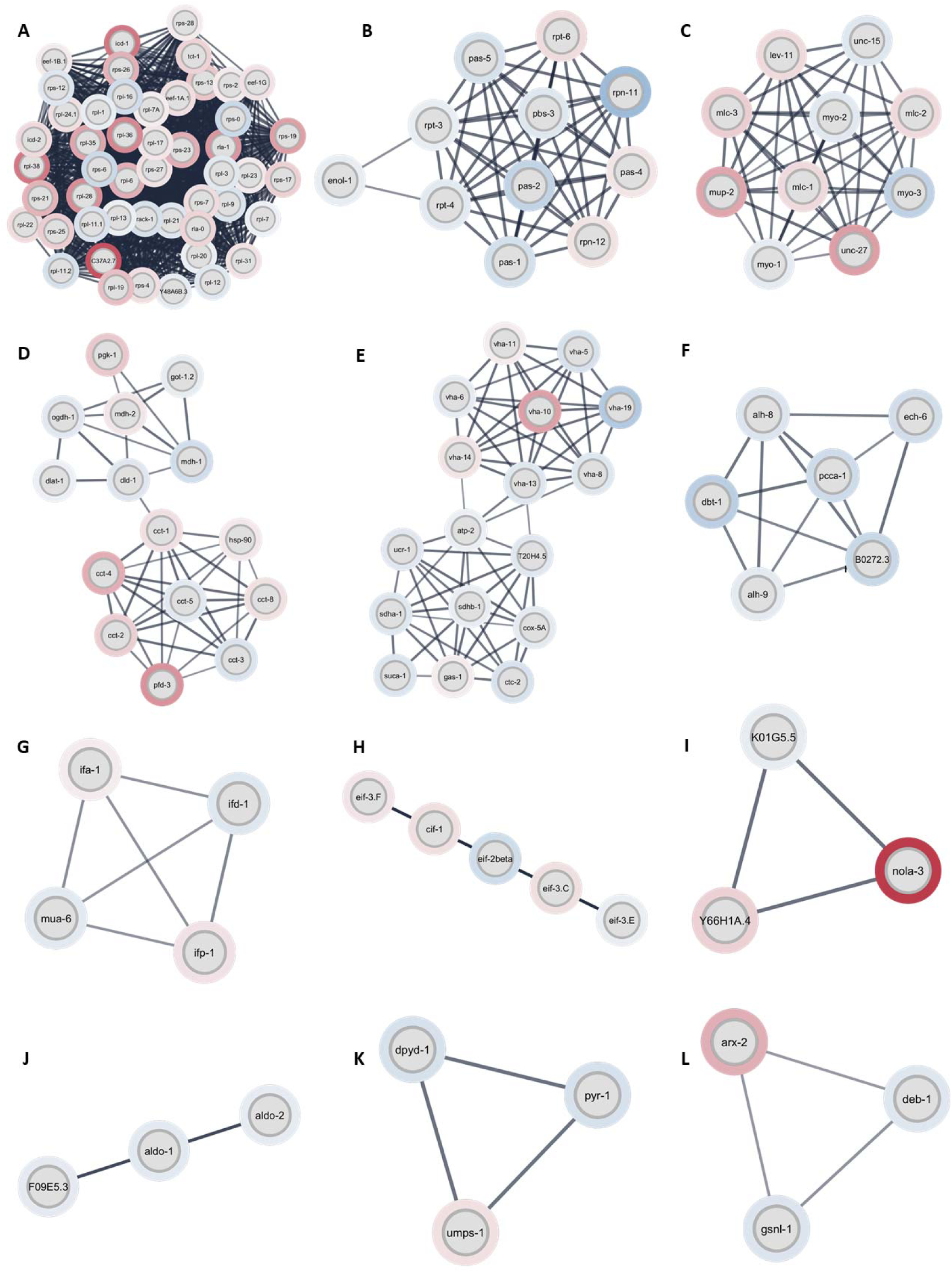
PPI networks from the N2 *C. elegans.* (A–L) Subnetworks were identified using the MCODE algorithm.

The largest module (Figure 6A) was strongly enriched in ribosomal proteins from both small and large subunits, together with translation elongation factors EEF-1A.1 and EEF-1G (Daftuar et al., 2013; B. Xu et al., 2022) and the scaffold protein RACK-1 (Kolosov et al., 2025). The high connectivity of these proteins identifies protein synthesis as a major functional hub within the CBZ-responsive network. This observation was further supported by Figure 6H, which contained translation initiation factors from the eIF3 complex and eIF2β, and by Figure 6I, in which NOLA-3 was a highly connected protein associated with RNA-related factors and ribosome biogenesis was a highly connected protein associated with RNA-related factors and ribosome biogenesis (Xu et al., 2022). Together, these modules indicate that CBZ affects multiple stages of protein synthesis, from ribosome biogenesis to translation initiation and elongation, consistent with the proteostasis alterations described above.

The proteasome-enriched module (Figure 6B), comprising PAS, PBS, RPN, and RPT subunits, further highlighted protein quality control as an important component of the response. Rather than indicating uniform activation, the coordinated alteration of these interacting proteins suggests remodeling of the ubiquitin–proteasome system, potentially supporting the removal of damaged or misfolded proteins (Peth et al., 2013). The pyrimidine metabolism module (Figure 6K), containing DPYD-1, PYR-1, and UMPS-1, additionally points to changes in nucleotide metabolism that may support cellular maintenance under stress.

Several modules were associated with mitochondrial and metabolic functions. Figure 6D contained proteins involved in glycolysis (PGK-1), the TCA cycle (MDH-1, MDH-2, OGDH-1), pyruvate metabolism (DLAT-1 and DID-1) (Penkov et al., 2020), and the chaperonin-containing TCP-1 complex (CCT proteins) (Leroux & Candido, 1997), whereas Figure 6E combined V-ATPase components with mitochondrial respiratory proteins, including SDHA-1, SDHB-1, ATP-2, and COX-5A (Song et al., 2020c). Figures 6J and 6F further represented glycolytic and lipid metabolic processes, respectively. The integration of these modules supports the coordinated remodeling of energy metabolism, mitochondrial function, lipid utilization, and protein folding, consistent with the metabolic adaptations identified in the proteomic analysis.

The network also revealed extensive structural remodeling. Figure 6C was enriched in muscle-associated proteins, including myosin (MYO-1, MYO-2, and MYO-3) (Johnson et al., 2022), myosin light chains (MLC-1, MLC-2, and MLC-3)(Rushforth et al., 1998), tropomyosin (LEV-11) (Kimmich et al., 2025), paramyosin (UNC-15) (Waterston et al., 1977), and troponins (UNC-27 and MUP-2) (Burkeen et al., 2004; Myers et al., 1996), while Figure 6G contained intermediate filament proteins, including IFA-1, IFB-1, IFD-1, and MUA-6 (Karabinos, 2019). Considering that CBZ exerts its fungicidal activity by disrupting microtubule polymerization (Winder, 2001), the modulation of proteins involved in muscle contraction and cytoskeletal organization suggests that similar structural disturbances occur in *C. elegans*. These alterations may contribute to impaired locomotion, muscle integrity, intracellular organization, and overall cellular architecture. Figure 6L included ANX-2, DEB-1, and GSN-1, which are involved in actin dynamics, membrane organization, and cell adhesion (Gerke et al., 2024; Hayes et al., 2004). Together, these modules indicate broad remodeling of the cytoskeletal and structural machinery, consistent with the established effects of CBZ on microtubule dynamics and potentially contributing to alterations in cellular architecture and locomotion.

In summary, the PPI analysis demonstrates that CBZ induces interconnected alterations across protein synthesis and proteostasis, mitochondrial and metabolic function, cytoskeletal organization, and intracellular processes, rather than affecting isolated proteins or pathways. The predominance of ribosomal and translation-related proteins among the highly connected components further supports protein synthesis as a central feature of the molecular response to CBZ.

## 4. Conclusion

This study provides an integrated toxicological assessment of CBZ and new mechanistic insights into its toxicity in *C. elegans*. By integrating quantitative proteomics with functional enrichment and protein–protein interaction network analyses, we demonstrate that CBZ toxicity extends beyond its classical microtubule-disrupting activity, inducing coordinated alterations in protein synthesis and proteostasis, mitochondrial function, energy and lipid metabolism, intracellular trafficking, chromatin organization, and redox homeostasis. Network and hub analyses further identified functionally connected protein modules, with ribosomal proteins and translation-associated factors emerging as major regulatory nodes, highlighting translational remodeling as a central component of the cellular response.

Overall, CBZ exposure elicited widespread molecular reprogramming despite subtle phenotypic effects, revealing substantial disturbances in cellular homeostasis following sublethal exposure. The evolutionary conservation and mammalian homology of several proteins identified here further support the translational relevance of these findings, providing mechanistic insights into cellular processes potentially affected by CBZ exposure in higher organisms, including humans. These results reinforce the value of *C. elegans* and integrative proteomics for elucidating conserved mechanisms of environmental toxicant action

## Acknowledgements/Funding

This project was supported by Conselho Nacional de Desenvolvimento Científico e Tecnológico (CNPq- grant #301808/2018-0 and scholarship for A.C.S), and INCT Nanotechnology for Sustainable Agriculture program (MCTI/CNPq-INCT NanoAgro, grant #405924/2022-4). We would like to thank Universidade Federal do Pampa and Universidade Federal de Pernambuco for funding and infrastructure, Fundação de Amparo à Pesquisa do Estado do Rio Grande do Sul (Grant #16/2551-0000248-7 and RITES Call grant # 22/2551-0000385-0), and Coordenação de Aperfeiçoamento de Pessoal de Nível Superior (CAPES, Finance Code 001). Authors would like to thank CGC for providing strains. CGC is funded by NIH Office of Research Infrastructure Programs (P40 OD010440). The authors acknowledge the use of artificial intelligence–based tools to assist with language editing and manuscript refinement. The authors take full responsibility for the content of the manuscript.

## Conflict of interest

The authors declare no conflict of interest.

## List of abbreviations

CBZ: carbendazim
NGM: nematode growth medium
DMSO: dimethylsulfoxide
DTT: dithiothreitol
USA: United States of America
LC: liquid chromatography
MS: mass
FDR: false discovery rate
GO: gene ontology
DAPs: differentially abundant proteins
BP: biological process
CC: cellular component
MF: molecular function
PPI: protein-protein interaction
FC: fold change
PCA: principal component analysis
CV: coefficient of variation
NAC: polypeptide-associated complex
UPR: unfolded protein response
ROS: reactive oxygen species
RNS: reactive nitrogen species
RNA: ribonucleic acid
DNA: deoxyribonucleic acid
ATP: adenosine triphosphate
TCA: tricarboxylic acid
aa: aminoacid
CC: closeness centrality
MCC: maximal clique centrality
SEM: mean ± standard error of the mean
IUPAC: International Union of Pure and Applied Chemistry

## Notes

https://github.com/joaoluna-dev/proteomics_c_elegans

https://drive.google.com/drive/folders/17RnPaCoLLtFzfssdO-ncgxyHWvDoXfUB

### Competing Interest Statement

The authors have declared no competing interest.

