## Supplementary material for "Carbendazim Disrupts Cellular Homeostasis and Proteostasis in *Caenorhabditis elegans*: A Global Proteomic Approach"

**Supplementary materials**

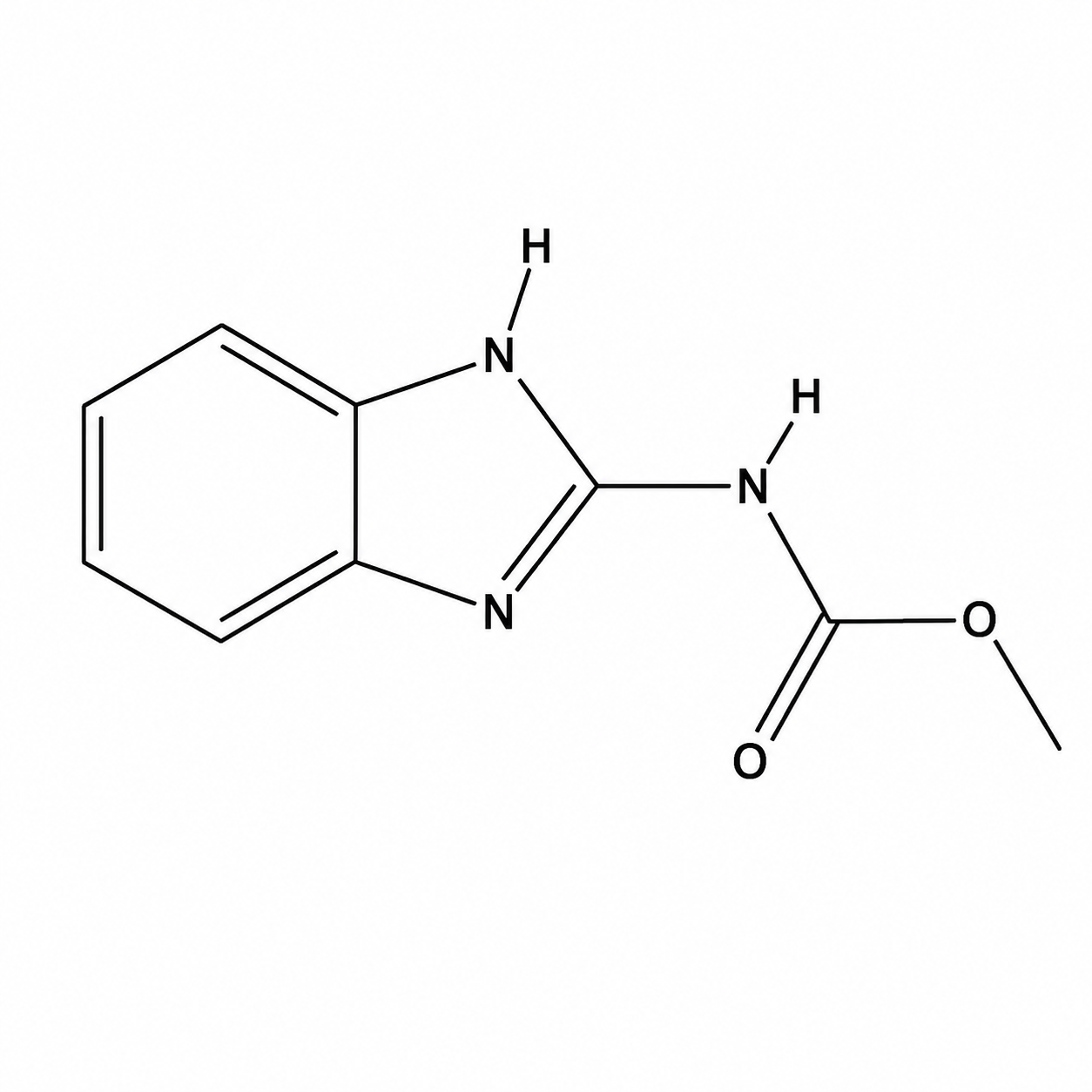

**Figure S1. Chemical structure of carbendazim (CBZ).** International Union of Pure and Applied Chemistry (IUPAC) name: methyl *N*-(1*H*-benzimidazol-2-yl)carbamate; molecular formula: [C_9_H_9_N_3_O_2_](https://pubchem.ncbi.nlm.nih.gov#query=C9H9N3O2); molecular weight: 191.19 g/mol. From PubChem (CID 25429).

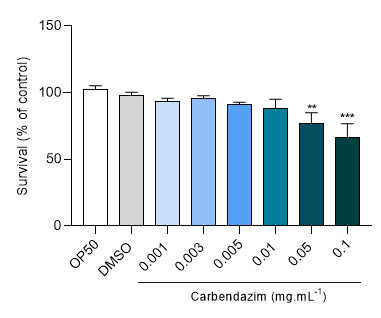

**Figure S2. Survival analysis.** (A) Survival rate (L4) of N2 *C. elegans* exposed to CBZ (n = 3–5). Data are expressed as the mean ± standard error of the mean (SEM). (*) Indicates a statistically significant difference from the control group: ***p < 0.01 and ***p < 0.001*, by one-way ANOVA followed by Tukey’s post hoc test.

**Table S1.** **Protein identification.** List of differentially abundant proteins (DAPs) of *C. elegans* against CBZ exposure (String Database, 2025)

| **protein_name** | **Accession** | **pAdjusted** | **-log10(pAdjusted)** | **log2(fc)** | **Information** |
| --- | --- | --- | --- | --- | --- |
| **Upregulated** | | | | | |
| ACBP-1 | O01805 | 0.002298507 | 2.638554147 | 5.340586076 | Acyl-CoA-binding protein homolog 1; Binds medium- and long-chain acyl-CoA esters with very high affinity and may function as an intracellular carrier of acyl-CoA esters. (86 aa) |
| NOLA-3 | Q9XVR8 | 0.006985594 | 2.155796682 | 4.500246699 | Putative H/ACA ribonucleoprotein complex subunit 3; Required for ribosome biogenesis. Part of a complex which catalyzes pseudouridylation of rRNA. This involves the isomerization of uridine such that the ribose is subsequently attached to C5, instead of the normal N1. Pseudouridine ('psi') residues may serve to stabilize the conformation of rRNAs (By similarity). (64 aa) |
| C37A2.7 | O01504 | 0.000250541 | 3.601120759 | 3.947049093 | 60S acidic ribosomal protein P2; Plays an important role in the elongation step of protein synthesis. (107 aa) |
| BAF-1 | Q03565 | 0.000534896 | 3.271730964 | 3.65661084 | Barrier-to-autointegration factor 1; DNA-binding protein which plays an essential role in nuclear envelope formation. Required for normal chromosome segregation during mitosis. Associates with the nuclear lamina via its interaction with LEM domain containing proteins emr-1 and lem-2. In association with lem-3, plays a role in radiation-induced DNA damage repair response. Belongs to the BAF family. |
| HIS-3 | P09588 | 8.00474E-05 | 4.096653001 | 3.331511226 | Histone H2A. |
| GBA-4 | Q9UB00 | 0.013242443 | 1.878031897 | 3.076311834 | Putative glucosylceramidase 4; Belongs to the glycosyl hydrolase 30 family. (519 aa) |
| ICD-1 | Q18885 | 0.000149859 | 3.824318129 | 3.048440257 | Transcription factor BTF3 homolog; May act as a transcription factor that exert a negative effect on the expression of several genes that are transcribed by RNA polymerase II. (161 aa) |
| RPL-38 | O17570 | 0.000278112 | 3.55577955 | 3.009011693 | 60S ribosomal protein L38; Belongs to the eukaryotic ribosomal protein eL38 family. (70 aa) |
| HIS-11 | P04255;Q27484;Q27876;Q27894 | 6.06261E-05 | 4.21734063 | 2.703868199 | Histone H2B 1. |
| RPL-36 | P49181 | 0.000819168 | 3.086626987 | 2.65213382 | 60S ribosomal protein L36; Component of the large ribosomal subunit. Belongs to the eukaryotic ribosomal protein eL36 family. |
| RPL-28 | Q21930 | 0.003355832 | 2.474199761 | 2.500856728 | 60S ribosomal protein L28; Belongs to the eukaryotic ribosomal protein eL28 family. (126 aa) |
| PFD-3 | O18054 | 0.001973368 | 2.704791838 | 2.400296467 | Probable prefoldin subunit 3; Binds specifically to cytosolic chaperonin (c-CPN) and transfers target proteins to it. Binds to nascent polypeptide chain and promotes folding in an environment in which there are many competing pathways for nonnative proteins (By similarity); Belongs to the prefoldin subunit alpha family. (185 aa) |
| LARP-1 | D5MCN2 | 0.000278112 | 3.55577955 | 2.268663884 | La-related protein 1; RNA-binding protein that promotes oogenesis by repressing fem-3 expression during germline development. Binds poly-U and poly-G stretches of RNA in vitro. Regulates target RNAs expression either by regulating their stability or translation. (1150 aa) |
| RPLP-1 | P91913 | 0.00017232 | 3.763663996 | 2.259519971 | RPL-11.1 is a large ribosomal subunit L11 protein in *Caenorhabditis elegans*, encoded by the autosomal gene *rpl-11.1* and primarily expressed in germline cells and pharyngeal muscle. Its human ortholog, ribosomal protein L11 (RPL-11), functions as a tumor suppressor through RPL11–MDM2 antagonism in the p53 pathway. |
| HSP-4 | P20163 | 0.000671189 | 3.173154892 | 2.159572088 | Endoplasmic reticulum chaperone BiP homolog; Endoplasmic reticulum chaperone that plays a key role in protein folding and quality control in the endoplasmic reticulum lumen (By similarity). Required for ER dynamics during the first embryonic cell divisions. Specifically, controls ER transition into sheet-like structures at the onset of mitosis, possibly by regulating homotypic membrane fusion. Belongs to the heat shock protein 70 family. (657 aa) |
| UNC-27 | Q9GYF1 | 6.05671E-05 | 4.217763365 | 2.108382482 | Troponin I 2; Troponin I is the inhibitory subunit of troponin, the thin filament regulatory complex which confers calcium-sensitivity to muscle actomyosin ATPase activity. (242 aa) |
| RPS-19 | O18650 | 4.01206E-05 | 4.396632378 | 2.082121339 | 40S ribosomal protein S19; Belongs to the eukaryotic ribosomal protein eS19 family. (146 aa) |
| VHA-10 | P91303 | 0.00097305 | 3.011864926 | 2.062219723 | Probable V-type proton ATPase subunit G; Catalytic subunit of the peripheral V1 complex of vacuolar ATPase (V-ATPase). V-ATPase is responsible for acidifying a variety of intracellular compartments in eukaryotic cells. (126 aa) |
| RPL-35 | P34662 | 0.006737792 | 2.171482379 | 1.902604575 | 60S ribosomal protein L35; Belongs to the universal ribosomal protein uL29 family. (123 aa) |
| MUP-2 | Q27371 | 0.001973368 | 2.704791838 | 1.902519151 | Troponin T; Troponin T is the tropomyosin-binding subunit of troponin, the thin filament regulatory complex which confers calcium-sensitivity to muscle actomyosin ATPase activity. (405 aa) |
| ADH-5 | Q17335 | 0.000139449 | 3.855583999 | 1.747816975 | Alcohol dehydrogenase class-3; Class-III ADH is remarkably ineffective in oxidizing ethanol, but it readily catalyzes the oxidation of long-chain primary alcohols and the oxidation of S-(hydroxymethyl) glutathione (By similarity). Plays a role in the calcium flux to the cytoplasm in the ASJ sensory neurons upon removal of a nitric oxide stimulus. (386 aa) |
| CCT-4 | P47208 | 0.002866231 | 2.542688763 | 1.746172418 | T-complex protein 1 subunit delta; Molecular chaperone; assists the folding of proteins upon ATP hydrolysis. Known to play a role, in vitro, in the folding of actin and tubulin. (540 aa) |
| RPL-6 | P47991 | 0.000163165 | 3.787372086 | 1.73188749 | 60S ribosomal protein L6; Component of the large ribosomal subunit. Belongs to the eukaryotic ribosomal protein eL6 family. (217 aa) |
| RPS-23 | Q19877 | 0.000163165 | 3.787372086 | 1.727326551 | 40S ribosomal protein S23; Belongs to the universal ribosomal protein uS12 family. (143 aa) |
| ARX-2 | P53489 | 0.000790486 | 3.102105648 | 1.709134773 | Actin-related protein 2; Functions as ATP-binding component of the Arp2/3 complex which is involved in regulation of actin polymerization and together with an activating nucleation-promoting factor (NPF) mediates the formation of branched actin networks. Seems to contact the pointed end of the daughter actin filament (By similarity). (395 aa) |
| F37C4.5 | O44400 | 4.01206E-05 | 4.396632378 | 1.653833988 | Protein F37C4.5. (556 aa) |
| RPS-26 | O45499 | 0.002937779 | 2.531980924 | 1.62918009 | 40S ribosomal protein S26; Belongs to the eukaryotic ribosomal protein eS26 family. (117 aa) |
| RPS-21 | P49197 | 0.00017232 | 3.763663996 | 1.620314431 | 40S ribosomal protein S21. (88 aa) |
| FAR-1 | P34382 | 0.000214153 | 3.669275519 | 1.609074889 | Fatty-acid and retinol-binding protein 1; Probably binds lipids; Belongs to the fatty-acid and retinol-binding protein (FARBP) family. (182 aa) |
| CYC-2.1 | P19974;Q23240 | 0.00025288 | 3.597085654 | 1.607463163 | Cytochrome c 2.1; Electron carrier protein. The oxidized form of the cytochrome c heme group can accept an electron from the heme group of the cytochrome c1 subunit of cytochrome reductase. Cytochrome c then transfers this electron to the cytochrome oxidase complex, the final protein carrier in the mitochondrial electron-transport chain. (111 aa) |
| ADS-1 | O45218 | 0.00039653 | 3.401724093 | 1.591082507 | Alkyldihydroxyacetonephosphate synthase; Catalyzes the exchange of an acyl for a long-chain alkyl group and the formation of the ether bond in the biosynthesis of ether phospholipids; Belongs to the FAD-binding oxidoreductase/transferase type 4 family. (597 aa) |
| CLIK-2 | Q09936 | 0.001109535 | 2.954859154 | 1.409054741 | Uncharacterized protein C53C9.2; Belongs to the calponin family. (397 aa) |
| LET-754 | P34346 | 0.021489234 | 1.667779062 | 1.376775574 | Adenylate kinase; Catalyzes the reversible transfer of the terminal phosphate group between ATP and AMP. Plays an important role in cellular energy homeostasis and in adenine nucleotide metabolism. Adenylate kinase activity is critical for regulation of the phosphate utilization and the AMP de novo biosynthesis pathways. (251 aa) |
| RPL-19 | O02639 | 0.00040735 | 3.39003235 | 1.329670973 | 60S ribosomal protein L19; Belongs to the eukaryotic ribosomal protein eL19 family. (198 aa) |
| FAR-2 | P34383 | 0.000317787 | 3.497863998 | 1.275758716 | Fatty-acid and retinol-binding protein 2; Probably binds lipids; Belongs to the fatty-acid and retinol-binding protein (FARBP) family. (182 aa) |
| HIS-1 | P62784 | 0.000923838 | 3.034403964 | 1.216128905 | Histone H4. |
| T19B4.3 | P91455 | 0.022283391 | 1.652018712 | 1.210746477 | Adenine phosphoribosyltransferase; Catalyzes a salvage reaction resulting in the formation of AMP, that is energically less costly than de novo synthesis. Belongs to the purine/pyrimidine phosphoribosyltransferase family. (185 aa) |
| DLC-1 | Q22799 | 0.013020324 | 1.885378225 | 1.210085017 | Dynein light chain 1, cytoplasmic; Acts as a non-catalytic accessory component of a dynein complex (By similarity). Part of a complex with bicd-1 and egal-1, which is recruited to the nuclear envelope by unc-83, where in turn, it recruits dynein to the nuclear surface and regulates nuclear migrations in hypodermal precursor cells. Probably within a dynein motor complex, plays a role in the cell fate specification of the germline and oogenesis. In particular, it inhibits germ cell proliferation. Regulates the function and localization of the RNA-binding protein fbf- 2 in the germline. |
| RPS-13 | P51404 | 0.000580012 | 3.236563228 | 1.183627908 | 40S ribosomal protein S13; Belongs to the universal ribosomal protein uS15 family. (151 aa) |
| PLP-1 | Q94230 | 0.042231207 | 1.374366503 | 1.131815377 | Pur alpha Like Protein. (226 aa) |
| PGK-1 | P91427 | 0.00010202 | 3.991315721 | 1.109518837 | Probable phosphoglycerate kinase; Belongs to the phosphoglycerate kinase family. (417 aa) |
| ADSL-1 | Q21774 | 0.040721706 | 1.390174041 | 1.103868487 | Adenylosuccinate lyase; Catalyzes two non-sequential steps in de novo AMP synthesis: converts (S)-2-(5-amino-1-(5-phospho-D-ribosyl)imidazole-4- carboxamido)succinate (SAICAR) to fumarate plus 5-amino-1-(5-phospho-D- ribosyl)imidazole-4-carboxamide, and thereby also contributes to de novo IMP synthesis, and converts succinyladenosine monophosphate (SAMP) to AMP and fumarate. (478 aa) |
| F37F2.2 | O61749 | 0.020780583 | 1.682342264 | 1.072546956 | Probable signal recognition particle 19 kDa protein; Signal-recogwago-1kin-3nition-particle assembly, binds directly to 7S RNA and mediates binding of the 54 kDa subunit of the SRP. (142 aa) |
| ACBP-3 | Q20507 | 0.006140128 | 2.211822579 | 1.054912139 | Acyl-CoA-binding protein homolog 3; Binds medium- and long-chain acyl-CoA esters with very high affinity and may function as an intracellular carrier of acyl-CoA esters. (116 aa) |
| WAGO-1 | Q21770 | 0.036535561 | 1.43728422 | 1.043623328 | Argonaute protein wago-1; Argonaute protein which is involved in the endogenous small interfering RNA (endo-siRNA) pathway. Interacts with secondary 22G- RNAs, which are RNA-dependent RNA polymerase-derived endo-siRNAs, typically 22 nucleotides in length with a 5'guanosine residue. In the germline, functions in a genome surveillance system to silence transposons and aberrant transcripts. |
| KIN-3 | P18334 | 0.000893314 | 3.048995621 | 1.043529196 | Casein kinase II subunit alpha; Casein kinases are operationally defined by their preferential utilization of acidic proteins such as caseins as substrates. The alpha chain contains the catalytic site. May participate in Wnt signaling. Modulates two aspects of male mating behavior; response to hermaphrodite contact and vulval location, acting in the same pathway as lov-1 and pkd-2. (360 aa) |
| SNR-6 | Q9XTU6 | 0.000720613 | 3.142297676 | 1.035633647 | Probable small nuclear ribonucleoprotein E; Associated with the spliceosome snRNP U1, U2, U4/U6 and U5. Belongs to the snRNP Sm proteins family. (90 aa) |
| RPS-25 | P52821 | 0.016600095 | 1.779889424 | 0.999511823 | 40S ribosomal protein S25; Belongs to the eukaryotic ribosomal protein eS25 family. (117 aa) |
| MLC-3 | P53014 | 0.006299023 | 2.200726818 | 0.973944878 | Myosin, essential light chain. (153 aa) |
| CCT-2 | P47207 | 0.000790417 | 3.102143812 | 0.965101184 | T-complex protein 1 subunit beta; Molecular chaperone; assists the folding of proteins upon ATP hydrolysis. Known to play a role, in vitro, in the folding of actin and tubulin; Belongs to the TCP-1 chaperonin family. (529 aa) |
| NKB-1 | Q93235 | 6.05671E-05 | 4.217763365 | 0.941308253 | Sodium/potassium-transporting ATPase subunit beta-1; Non-catalytic component of the active enzyme, which catalyzes the hydrolysis of ATP coupled with the exchange of Na(+) and K(+) ions across the plasma membrane. The beta subunit regulates, through assembly of alpha/beta heterodimers, the number of sodium pumps transported to the plasma membrane (By similarity). May also may function in the peptidic vesicle transport pathway from the endoplasmic reticulum for the defecation behaviors. (320 aa) |
| RPS-17 | O01692 | 0.021273546 | 1.672160105 | 0.920886005 | 40S ribosomal protein S17; Belongs to the eukaryotic ribosomal protein eS17 family. (130 aa) |
| HUM-6 | P91443 | 0.038010111 | 1.420100858 | 0.898054481 | Unconventional myosin heavy chain 6; Myosins are actin-based motor molecules with ATPase activity. Unconventional myosins serve in intracellular movements. Their highly divergent tails are presumed to bind to membranous compartments, which would be moved relative to actin filaments (By similarity). (2098 aa) |
| GARR-1 | Q9TYK1 | 0.000815301 | 3.088681998 | 0.893538287 | GARR-1 (GAR1 related) is a nucleolar protein in *C. elegans* identified through a survey of RG/RGG repeat-containing proteins, sharing homology with the conserved nucleolar protein GAR1. The protein was named GARR-1 based on predicted shared homology with GAR1 detected through HHpred and MARRVEL toolkits It accumulates in germ cell nucleoli throughout all stages of worm development, displaying a "net-like" sub-nucleolar structure similar to the nucleolar protein FIB-1. |
| PAR-5 | P41932 | 0.000163165 | 3.787372086 | 0.861688195 | 14-3-3-like protein 1. |
| TCT-1 | Q93573 | 0.000790486 | 3.102105648 | 0.843795985 | Translationally-controlled tumor protein homolog; Involved in calcium binding and microtubule stabilization. Belongs to the TCTP family. (181 aa) |
| PFN-2 | Q20025 | 0.007912718 | 2.10167432 | 0.84277308 | Profilin-2; Binds to actin and affects the structure of the cytoskeleton. At high concentrations, profilin prevents the polymerization of actin, whereas it enhances it at low concentrations. By binding to PIP2, it inhibits the formation of IP3 and DG (By similarity). (131 aa) |
| RLA-0 | Q93572 | 0.001050382 | 2.978652592 | 0.838271011 | 60S acidic ribosomal protein P0; Ribosomal protein P0 is the functional equivalent of E.coli protein L10. (312 aa) |
| KIN-19 | P42168 | 0.001375707 | 2.861474206 | 0.798190781 | Casein kinase I isoform alpha; Belongs to the protein kinase superfamily. CK1 Ser/Thr protein kinase family. Casein kinase I subfamily. (341 aa) |
| UNC-87 | P37806 | 0.000965135 | 3.015411905 | 0.787159424 | Protein unc-87; Thin filament-associated protein that is implicated in actin bundling and actin filament dynamics. Exhibits F-actin cross-linking activity. Required for the maintenance of sarcomeric actin organization in striated muscles. Competes with unc-60 isoform b for actin binding and protects actin filaments from depolymerization by unc-60, thereby contributing to actin filament stability. Cooperates with myosin to form actomyosin bundles and inhibits actomyosin ATPase activity and actomyosin motility. Might protect the myofilaments from mechanical stress. |
| LONP-2 | Q9XW87 | 0.001250055 | 2.903070743 | 0.767899291 | Lon protease homolog 2, peroxisomal; ATP-dependent serine protease that mediates the selective degradation of misfolded and unassembled polypeptides in the peroxisomal matrix. Necessary for type 2 peroxisome targeting signal (PTS2)-containing protein processing and facilitates peroxisome matrix protein import. (773 aa) |
| CLU-1 | P34466 | 0.001368615 | 2.863718661 | 0.705766739 | Clustered mitochondria protein homolog; mRNA-binding protein involved in proper cytoplasmic distribution of mitochondria; Belongs to the CLU family. (1247 aa) |
| ICD-2 | Q86S66 | 0.001161534 | 2.934968168 | 0.697011294 | Nascent polypeptide-associated complex subunit alpha; May promote appropriate targeting of ribosome-nascent polypeptide complexes. (197 aa) |
| F38B2.4 | Q20140 | 0.003405471 | 2.46782286 | 0.688476426 | Adenylate kinase isoenzyme 1; Catalyzes the reversible transfer of the terminal phosphate group between ATP and AMP. Plays an important role in cellular energy homeostasis and in adenine nucleotide metabolism. (210 aa) |
| LEV-11 | Q27249 | 0.001653142 | 2.781689885 | 0.684599904 | Tropomyosin isoforms a/b/d/f; Tropomyosin, in association with the troponin complex, plays a central role in the calcium dependent regulation of muscle contraction. Involved in muscle actin filament organization and muscle arm extension and morphology. Protects actin filaments from depolymerization by unc-60 in vitro. Also has a role in male mating behavior by regulating the copulatory spicules. Binds to F-actin. |
| MLC-1 | P19625 | 0.047040218 | 1.327530672 | 0.683152836 | Myosin regulatory light chain 1. (170 aa) |
| MLC-2 | P19626 | 0.047040218 | 1.327530672 | 0.683152836 | Myosin regulatory light chain 2. |
| RPL-22 | P52819 | 0.00361326 | 2.442100796 | 0.667758425 | 60S ribosomal protein L22. (130 aa) |
| RPT-6 | Q9XTT9 | 0.005406058 | 2.267119292 | 0.661243316 | 26S proteasome regulatory subunit 8; Component of the 26S proteasome, a multiprotein complex involved in the ATP-dependent degradation of ubiquitinated proteins. This complex plays a key role in the maintenance of protein homeostasis by removing misfolded or damaged proteins, which could impair cellular functions, and by removing proteins whose functions are no longer required. Therefore, the proteasome participates in numerous cellular processes, including cell cycle progression, apoptosis, or DNA damage repair (By similarity). Belongs to the heterohexameric ring of AAA (ATPases Associated with diverse cellular Activities). |
| AHCY-1 | P27604 | 0.000790486 | 3.102105648 | 0.637386805 | Adenosylhomocysteinase; Adenosylhomocysteine is a competitive inhibitor of S- adenosyl-L-methionine-dependent methyl transferase reactions; therefore adenosylhomocysteinase may play a key role in the control of methylations via regulation of the intracellular concentration of adenosylhomocysteine. (437 aa) |
| RPS-7 | Q23312 | 0.000720613 | 3.142297676 | 0.631711212 | 40S ribosomal protein S7; Belongs to the eukaryotic ribosomal protein eS7 family. (194 aa) |
| RPS-4 | Q9N3X2 | 0.001050382 | 2.978652592 | 0.618164851 | 40S ribosomal protein S4; Belongs to the eukaryotic ribosomal protein eS4 family. (259 aa) |
| FAT-2 | G5EGA5 | 0.006985594 | 2.155796682 | 0.604371389 | Delta(12) fatty acid desaturase fat-2; Catalyzes the desaturation of oleic acid (Delta(9)-18:1, or C18:1(n-9)) to linoleic acid (Delta(9,12)-18:2, or C18:2(n-6)). This desaturation event is essential for the synthesis of longer chain polyunsaturated fatty acids (PUFAs) such as arachidonic acid (AA, or C20:4(n-6)) and eicosapentaenoic acid (EPA, or C20:5(n-3)). |
| EFT-3 | P53013 | 0.004544595 | 2.342504817 | 0.594912741 | Elongation factor 1-alpha; This protein promotes the GTP-dependent binding of aminoacyl- tRNA to the A-site of ribosomes during protein biosynthesis; Belongs to the TRAFAC class translation factor GTPase superfamily. Classic translation factor GTPase family. EF-Tu/EF-1A subfamily. (463 aa) |
| EIF-3.C | O02328 | 0.02155893 | 1.666372802 | 0.574378957 | Eukaryotic translation initiation factor 3 subunit C; Component of the eukaryotic translation initiation factor 3 (eIF-3) complex, which is involved in protein synthesis of a specialized repertoire of mRNAs and, together with other initiation factors, stimulates binding of mRNA and methionyl-tRNAi to the 40S ribosome. The eIF-3 complex specifically targets and initiates translation of a subset of mRNAs involved in cell proliferation. (898 aa) |
| RPL-24.1 | O01868 | 0.001061146 | 2.974224733 | 0.568481518 | 60S ribosomal protein L24; Belongs to the eukaryotic ribosomal protein eL24 family. (159 aa) |
| CCT-1 | P41988 | 0.000911101 | 3.040433494 | 0.568263628 | T-complex protein 1 subunit alpha; Molecular chaperone; assists the folding of proteins upon ATP hydrolysis. Known to play a role, in vitro, in the folding of actin and tubulin. (549 aa) |
| LBP-2 | Q20224;Q20223 | 0.002298507 | 2.638554147 | 0.566895324 | Fatty acid-binding protein homolog 2; May play a role in sequestering potentially toxic fatty acids and their peroxidation products, or it may be involved in the maintenance of the impermeable lipid layer of the eggshell. (161 aa) |
| CIF-1 | Q94261 | 0.016072803 | 1.793908391 | 0.564480208 | COP9/Signalosome and eIF3 complex-shared subunit 1; Component of the eukaryotic translation initiation factor 3 (eIF-3) complex, which is involved in protein synthesis of a specialized repertoire of mRNAs and, together with other initiation factors, stimulates binding of mRNA and methionyl-tRNAi to the 40S ribosome. The eIF-3 complex specifically targets and initiates translation of a subset of mRNAs involved in cell proliferation (Potential). Component of the COP9 signalosome complex (CSN), a complex involved in various cellular and developmental processes. |
| HTZ-1 | Q27511 | 0.012444186 | 1.905033515 | 0.557738498 | Histone H2A.V; Variant histone H2A which replaces conventional H2A in a subset of nucleosomes. Nucleosomes wrap and compact DNA into chromatin, limiting DNA accessibility to the cellular machineries which require DNA as a template. Histones thereby play a central role in transcription regulation, DNA repair, DNA replication and chromosomal stability. DNA accessibility is regulated via a complex set of post- translational modifications of histones, also called histone code, and nucleosome remodeling (By similarity). Required to maintain non-distal tip cell (DTC) fate of somatic gonadal cells. |
| PMT-1 | Q23552 | 0.011270652 | 1.948050948 | 0.551877621 | Methyltransf_11 domain-containing protein. (484 aa) |
| CCT-8 | Q9N358 | 0.004780822 | 2.320497387 | 0.551404467 | T-complex protein 1 subunit theta; Molecular chaperone; assists the folding of proteins upon ATP hydrolysis. Known to play a role, in vitro, in the folding of actin and tubulin (By similarity). Required for correct subcellular localization of pgl-1; Belongs to the TCP-1 chaperonin family. (548 aa) |
| UMPS-1 | G5EDZ2 | 0.001368615 | 2.863718661 | 0.541321063 | Orotidine 5'-phosphate decarboxylase; Bifunctional enzyme which catalyzes the formation of UMP from orotate in the de novo pathway of pyrimidine biosynthesis. May also form UMP from uracil. Regulates the size of gut granules during embryonic development. Involved in resistance to DNA damaging agents including UV-C and X-ray radiation. In the C-terminal section; belongs to the OMP decarboxylase family. (497 aa) |
| RPS-27 | Q9TXP0 | 0.039572725 | 1.402604048 | 0.52537758 | 40S ribosomal protein S27. (83 aa) |
| RPL-31 | Q9U332 | 0.038685752 | 1.412448955 | 0.512737093 | 60S ribosomal protein L31. (122 aa) |
| IDHB-1 | Q93353 | 0.008894539 | 2.050876535 | 0.508521081 | Probable isocitrate dehydrogenase [NAD] subunit beta, mitochondrial; Belongs to the isocitrate and isopropylmalate dehydrogenases family. (379 aa) |
| FARS-3 | Q19713 | 0.0164888 | 1.782810949 | 0.498462594 | Phenylalanine--tRNA ligase beta subunit. (591 aa) |
| GLP-4 | Q9U1Q4 | 0.001103621 | 2.957179902 | 0.494368611 | Valine--tRNA ligase. |
| IFP-1 | Q09501 | 0.01412265 | 1.850083812 | 0.492242822 | Intermediate filament protein ifp-1; Cytoplasmic intermediate filaments provide mechanical strength to cells. Not essential protein. (776 aa) |
| EEF-1G | P54412 | 0.010760425 | 1.968170579 | 0.484602954 | Probable elongation factor 1-gamma; Probably plays a role in anchoring the complex to other cellular components. (398 aa) |
| CYN-5 | P52013;P52014 | 0.021769709 | 1.662147377 | 0.478899413 | Peptidyl-prolyl cis-trans isomerase 5; PPIases accelerate the folding of proteins. It catalyzes the cis-trans isomerization of proline imidic peptide bonds in oligopeptides. (204 aa) |
| GSY-1 | Q9U2D9 | 0.004373503 | 2.359170574 | 0.476070803 | Glycogen [starch] synthase; Transfers the glycosyl residue from UDP-Glc to the non- reducing end of alpha-1,4-glucan; Belongs to the glycosyltransferase 3 family. |
| RPN-12 | Q23449 | 0.013020324 | 1.885378225 | 0.475849017 | 26S proteasome non-ATPase regulatory subunit 8; Acts as a regulatory subunit of the 26S proteasome which is involved in the ATP-dependent degradation of ubiquitinated proteins. (250 aa) |
| TTR-2 | P34500 | 0.010608697 | 1.974337939 | 0.456949641 | Transthyretin-like protein 2. (148 aa) |
| VHA-14 | P34462 | 0.028409888 | 1.546530478 | 0.453498353 | V-type proton ATPase subunit D; Subunit of the peripheral V1 complex of vacuolar ATPase. V- ATPase is responsible for acidifying a variety of intracellular compartments in eukaryotic cells, thus providing most of the energy required for transport processes in the vacuolar system; Belongs to the V-ATPase D subunit family. (257 aa) |
| HSP-1 | P09446 | 0.000893314 | 3.048995621 | 0.449851315 | Heat shock 70 kDa protein A. (640 aa) |
| GST-36 | Q09607 | 0.011817722 | 1.927466215 | 0.43760778 | Probable glutathione S-transferase gst-36; Conjugation of reduced glutathione to a wide number of exogenous and endogenous hydrophobic electrophiles; Belongs to the GST superfamily. Sigma family. (210 aa) |
| EIF-3.F | Q18967 | 0.033889317 | 1.46993719 | 0.433320006 | Eukaryotic translation initiation factor 3 subunit F; Component of the eukaryotic translation initiation factor 3 (eIF-3) complex, which is involved in protein synthesis of a specialized repertoire of mRNAs and, together with other initiation factors, stimulates binding of mRNA and methionyl-tRNAi to the 40S ribosome. The eIF-3 complex specifically targets and initiates translation of a subset of mRNAs involved in cell proliferation. (294 aa) |
| UBC-18 | Q21633 | 0.006789496 | 2.168162456 | 0.430214998 | UBIQUITIN_CONJUGAT_2 domain-containing protein. |
| TBB-2 | P52275 | 0.002076977 | 2.682568297 | 0.427820933 | Tubulin beta-2 chain; Tubulin is the major constituent of microtubules. It binds two moles of GTP, one at an exchangeable site on the beta chain and one at a non-exchangeable site on the alpha chain; Belongs to the tubulin family. (450 aa) |
| RPL-17 | Q9BL19 | 0.001109535 | 2.954859154 | 0.427345241 | 60S ribosomal protein L17. (187 aa) |
| PAS-4 | Q95005 | 0.002298507 | 2.638554147 | 0.420694547 | Proteasome subunit alpha type-7; The proteasome is a multicatalytic proteinase complex which is characterized by its ability to cleave peptides with Arg, Phe, Tyr, Leu, and Glu adjacent to the leaving group at neutral or slightly basic pH. The proteasome has an ATP-dependent proteolytic activity. (253 aa) |
| GST-7 | P91253;P91254 | 0.010952062 | 1.9605041 | 0.420339461 | Probable glutathione S-transferase 7; Conjugation of reduced glutathione to a wide number of exogenous and endogenous hydrophobic electrophiles (By similarity). May play a role in the detoxification of reactive oxygen species produced during pathogenic bacterial infection. Belongs to the GST superfamily. Sigma family. (206 aa) |
| EIF-6 | O62106 | 0.026103189 | 1.583306437 | 0.419532399 | Eukaryotic translation initiation factor 6; Binds to the 60S ribosomal subunit and prevents its association with the 40S ribosomal subunit to form the 80S initiation complex in the cytoplasm. May also be involved in ribosome biogenesis. Involved in miRNA-mediated gene silencing. (246 aa) |
| RAN-1 | O17915 | 0.030903637 | 1.509990412 | 0.410530179 | GTP-binding nuclear protein ran-1; Ran GTPase system comprises ran-1, ran-2 and ran-3 and is essential in nucleocytoplasmic transport. Ran-1 is a GTP-binding protein that mediates the interaction between mitotic chromosomes and kinetochore microtubules. Plays a crucial role in nuclear envelope assembly at the end of each cell division. Required for the import of protein into the nucleus and also for RNA export. RCC1 (ran-3)/Ran (ran-1) complex (together with other proteins) acts as a component of a signal transmission pathway that detects unreplicated DNA. |
| CTH-2 | P55216 | 0.000498301 | 3.302508538 | 0.404120212 | Putative cystathionine gamma-lyase 2; Belongs to the trans-sulfuration enzymes family. (392 aa) |
| OST-1 | P34714 | 0.009063563 | 2.042701053 | 0.400704788 | SPARC; Has a high affinity for collagen. Affects nematode body morphology and mobility. Essential for *C. elegans* development and muscle function. The cysteine-rich region could have protease inhibitory activity or may provide the framework for a protein binding module. Probable role in skeletal morphogenesis; Belongs to the SPARC family. |
| MDH-2 | O02640 | 0.026291426 | 1.580185861 | 0.399316369 | Probable malate dehydrogenase, mitochondrial. (341 aa) |
| SAR-1 | Q23445 | 0.006985594 | 2.155796682 | 0.388695422 | GTP-binding protein SAR1; Involved in transport from the endoplasmic reticulum to the Golgi apparatus; Belongs to the small GTPase superfamily. SAR1 family. (193 aa) |
| ASB-1 | Q20053 | 0.021630829 | 1.664926834 | 0.380186901 | ATP Synthase B homolog. (301 aa) |
| CAP-2 | P34686 | 0.001432479 | 2.843911861 | 0.369555617 | F-actin-capping protein subunit beta; F-actin-capping proteins bind in a Ca(2+)-independent manner to the fast growing ends of actin filaments (barbed end) thereby blocking the exchange of subunits at these ends. Unlike other capping proteins (such as gelsolin and severin), these proteins do not sever actin filaments. |
| HADB-1 | P34255 | 0.026140048 | 1.582693615 | 0.345622861 | Predicted to enable acetyl-CoA C-acetyltransferase activity. Predicted to be involved in fatty acid beta-oxidation. Located in mitochondrion. Is expressed in body wall musculature; hypodermis; intestine; and tail. Human ortholog(s) of this gene implicated in mitochondrial metabolism disease and mitochondrial trifunctional protein deficiency 2. Orthologous to human HADHB (hydroxyacyl-CoA dehydrogenase trifunctional multienzyme complex subunit beta). |
| HSP-90 | Q18688 | 0.049468332 | 1.305672733 | 0.343652343 | Heat shock protein 90; Molecular chaperone that promotes the maturation, structural maintenance and proper regulation of specific target proteins involved for instance in cell cycle control and signal transduction. Undergoes a functional cycle that is linked to its ATPase activity. This cycle probably induces conformational changes in the client proteins, thereby causing their activation. Interacts dynamically with various co- chaperones that modulate its substrate recognition, ATPase cycle and chaperone function. By stabilizing the receptor-type guanylate cyclase daf-11 or another sig [...] (702 aa) |
| RPL-23 | P48158 | 0.016089823 | 1.793448746 | 0.312916643 | 60S ribosomal protein L23; Belongs to the universal ribosomal protein uL14 family. (140 aa) |
| F32D1.5 | O16294 | 0.003646688 | 2.438101345 | 0.309596982 | GMP reductase; Catalyzes the irreversible NADPH-dependent deamination of GMP to IMP. It functions in the conversion of nucleobase, nucleoside and nucleotide derivatives of G to A nucleotides, and in maintaining the intracellular balance of A and G nucleotides. (358 aa) |
| RPS-28 | Q95Y04 | 0.023846332 | 1.622578417 | 0.304200908 | 40S ribosomal protein S28; Belongs to the eukaryotic ribosomal protein eS28 family. (65 aa) |
| RPS-2 | P51403 | 0.013020324 | 1.885378225 | 0.302364764 | 40S ribosomal protein S2; Belongs to the universal ribosomal protein uS5 family. (272 aa) |
| EEF-1B.1 | P34460 | 0.014109454 | 1.850489798 | 0.29784562 | Probable elongation factor 1-beta/1-delta 1; EF-1-beta and EF-1-delta stimulate the exchange of GDP bound to EF-1-alpha to GTP; Belongs to the EF-1-beta/EF-1-delta family. (213 aa) |
| LET-721 | Q11190 | 0.00250321 | 2.601502785 | 0.288354917 | Electron transfer flavoprotein-ubiquinone oxidoreductase, mitochondrial; Accepts electrons from ETF and reduces ubiquinone. Belongs to the ETF-QO/FixC family. (597 aa) |
| GAS-1 | Q93873 | 0.034521743 | 1.46190729 | 0.280240682 | Probable NADH dehydrogenase [ubiquinone] iron-sulfur protein 2, mitochondrial; Core subunit of the mitochondrial membrane respiratory chain NADH dehydrogenase (Complex I) that is believed to belong to the minimal assembly required for catalysis. Complex I functions in the transfer of electrons from NADH to the respiratory chain. The immediate electron acceptor for the enzyme is believed to be ubiquinone (By similarity). (482 aa) |
| IFA-1 | P90901 | 0.018169168 | 1.740664965 | 0.27336204 | Intermediate filament protein ifa-1; Cytoplasmic intermediate filaments make up the structural component of the cytoskeleton providing mechanical strength to cells. Essential protein required during embryogenesis especially for survival past the L1 larva stage, involved in intestine morphogenesis. |
| VHA-11 | Q9XXU9 | 0.005776309 | 2.238349566 | 0.272793958 | V-type proton ATPase subunit C; Subunit of the peripheral V1 complex of vacuolar ATPase. Subunit C is necessary for the assembly of the catalytic sector of the enzyme and is likely to have a specific function in its catalytic activity. V-ATPase is responsible for acidifying a variety of intracellular compartments in eukaryotic cells. Has roles in embryogenesis and ovulation. (384 aa) |
| RPL-7A | Q966C6 | 0.029469175 | 1.530632028 | 0.270249393 | 60S ribosomal protein L7a; Belongs to the eukaryotic ribosomal protein eL8 family. (265 aa) |
| **Downregulated** | | | | | |
| IFA-3 | Q21065 | 0.037542525 | 1.425476517 | -0.263079721 | Intermediate filament protein ifa-3; Cytoplasmic intermediate filaments provide mechanical strength to cells. Essential protein, involved in attachment structures in epidermal cells that connect muscles to the external cuticle. Required for epidermal morphogenesis in embryos. Probable component of embryonic epidermal attachment structures. (581 aa) |
| RPL-7 | O01802 | 0.006985594 | 2.155796682 | -0.264157752 | 60S ribosomal protein L7; Binds to G-rich structures in 28S rRNA and in mRNAs. Plays a regulatory role in the translation apparatus; inhibits cell-free translation of mRNAs (By similarity). (244 aa) |
| PFN-1 | Q9XW16 | 0.039572725 | 1.402604048 | -0.266740276 | Profilin-1; Binds to actin and affects the structure of the cytoskeleton. At high concentrations, profilin prevents the polymerization of actin, whereas it enhances it at low concentrations. By binding to PIP2, it inhibits the formation of IP3 and DG. Also binds to poly(L-proline) and phosphatidylinositol 4,5-bisphosphate micelles. (132 aa) |
| DLAT-1 | Q19749 | 0.017429614 | 1.758712239 | -0.269701336 | Pyruvate dehydrogenase E2 component (dihydrolipoamide acetyltransferase); The pyruvate dehydrogenase complex catalyzes the overall conversion of pyruvate to acetyl-CoA and CO(2). It contains multiple copies of three enzymatic components: pyruvate dehydrogenase (E1), dihydrolipoamide acetyltransferase (E2) and lipoamide dehydrogenase (E3). (507 aa) |
| SKR-1 | G5ECU1 | 0.043297899 | 1.363533177 | -0.275043303 | Skp1-related protein; Probable essential component of SCF (SKP1-CUL1-F-box protein) E3 ubiquitin-protein ligase complexes, which mediate the ubiquitination and subsequent proteasomal degradation of target proteins. Regulates cell proliferation during embryonic and larval development. Involved in synapse elimination in early synapse development. May negatively regulate the apoptotic activity of cep-1 in response to genotoxic stress. Plays a role in sex determination. (176 aa) |
| APY-1 | Q19202 | 0.027540401 | 1.560029735 | -0.276412823 | Apyrase apy-1; Hydrolyzes UDP and to a lesser extent GDP. By preventing the accumulation of NDP, may promote the reglucosylation of incompletely folded glycoproteins in the endoplasmic reticulum following the unfolded protein response. |
| RPL-18A | O44480 | 0.028409888 | 1.546530478 | -0.276440024 | RPL-20 is the *C. elegans* ortholog of mammalian RPL18a/eL20, a component of the 60S large ribosomal subunit. A gain-of-function missense mutation in *rpl-20* suppresses gonadal cell migration defects caused by loss of the ADAMTS metalloprotease MIG-17. |
| ATP-2 | P46561 | 0.019998214 | 1.699008794 | -0.278287955 | ATP synthase subunit beta, mitochondrial; Mitochondrial membrane ATP synthase (F(1)F(0) ATP synthase or Complex V) produces ATP from ADP in the presence of a proton gradient across the membrane which is generated by electron transport complexes of the respiratory chain. F-type ATPases consist of two structural domains, F(1) - containing the extramembraneous catalytic core, and F(0) - containing the membrane proton channel, linked together by a central stalk and a peripheral stalk. |
| ATAD-3 | Q20748 | 0.016600095 | 1.779889424 | -0.288573682 | ATPase family AAA domain-containing protein 3; Essential for mitochondrial network organization, mitochondrial metabolism and cell growth at organism and cellular level. Important during development for the up-regulation of mitochondrial activity during the transition to higher larval stages. Regulates mitochondrial iron homeostasis. May play an important role in mitochondrial protein synthesis (By similarity). May also participate in mitochondrial DNA replication (By similarity). May bind to mitochondrial DNA D-loops and contribute to nucleoid stability (By similarity). (595 aa) |
| RPL-13 | P91128 | 0.030541339 | 1.515111931 | -0.291857831 | 60S ribosomal protein L13. (207 aa) |
| ICL-1 | Q10663 | 0.010952062 | 1.9605041 | -0.296578202 | Bifunctional glyoxylate cycle protein; In the C-terminal section; belongs to the malate synthase family. (968 aa) |
| ARF-1 | Q10943 | 0.005643979 | 2.24841463 | -0.301279746 | ADP-ribosylation factor 1-like 2; GTP-binding protein involved in protein trafficking; may modulate vesicle budding and uncoating within the Golgi apparatus. Involved in endoplasmic reticulum dynamics during embryogenesis. Also required for adult germline function. Plays a role in cell shedding during embryogenesis probably by promoting the endocytosis of cell adhesion molecules. During neurogenesis, involved in cell autonomous Q.p neuroblast asymmetric divisions that generate one precursor cell and one apoptotic cell, probably by controlling endocytosis. *[a.k.a. B0336.2, ADP-ribosylation factor 1-like 2, CCD61516.1, arf-1]* |
| EIF-3.E | O61820 | 0.002937779 | 2.531980924 | -0.318907823 | Eukaryotic translation initiation factor 3 subunit E; Component of the eukaryotic translation initiation factor 3 (eIF-3) complex, which is involved in protein synthesis of a specialized repertoire of mRNAs and, together with other initiation factors, stimulates binding of mRNA and methionyl-tRNAi to the 40S ribosome. The eIF-3 complex specifically targets and initiates translation of a subset of mRNAs involved in cell proliferation. (432 aa) |
| K12H4.7 | P34528 | 6.05671E-05 | 4.217763365 | -0.324542579 | Putative serine protease K12H4.7. (510 aa) |
| F40A3.3 | O16264 | 0.026314163 | 1.579810443 | -0.328840671 | Phosphatidylethanolamine-binding protein homolog F40A3.3. (226 aa) |
| LBP-6 | O01812 | 0.01844195 | 1.734193151 | -0.341997129 | Fatty acid-binding protein homolog 6. (135 aa) |
| SDHB-1 | Q09545 | 0.015081127 | 1.821566197 | -0.344600894 | Succinate dehydrogenase [ubiquinone] iron-sulfur subunit, mitochondrial; Iron-sulfur protein (IP) subunit of succinate dehydrogenase (SDH) that is involved in complex II of the mitochondrial electron transport chain and is responsible for transferring electrons from succinate to ubiquinone (coenzyme Q). (298 aa) |
| ENOL-1 | Q27527 | 0.010760425 | 1.968170579 | -0.350545213 | Enolase; Belongs to the enolase family. |
| RPT-3 | P46502 | 0.029989705 | 1.523027809 | -0.353894488 | Probable 26S proteasome regulatory subunit 6B; The 26S proteasome is involved in the ATP-dependent degradation of ubiquitinated proteins. The regulatory (or ATPase) complex confers ATP dependency and substrate specificity to the 26S complex (By similarity). |
| DHRS-4 | G5EGA6 | 0.007037511 | 2.152580899 | -0.354766607 | Dehydrogenase/reductase SDR family member 4; Catalyzes the reduction of isatin, 4-oxonon-2-enal, 9,10- phenanthrenequinone, menadione, 2,3-hexaenadione, 3,4-hexanedione and 2,3-heptanedione. (260 aa) |
| RPL-1 | Q9N4I4 | 0.005754093 | 2.240023152 | -0.358229146 | 60S ribosomal protein L10a; Belongs to the universal ribosomal protein uL1 family. (216 aa) |
| CDC-48.1 | P54811 | 0.017429614 | 1.758712239 | -0.358337518 | Transitional endoplasmic reticulum ATPase homolog 1; ATP-dependent chaperone which probably uses the energy provided by ATP hydrolysis to generate mechanical force to unfold substrate proteins, disassemble protein complexes, and disaggregate protein aggregates. Can also prevent aggregation of unfolded proteins also in an ATP- independent manner. Targets polyubiquitinated proteins for proteasomal degradation by binding to 'Lys-48'-linked polyubiquitin chains. Involved in the cytoplasmic elimination of misfolded proteins exported from the ER. |
| APP-1 | O44750 | 0.002298507 | 2.638554147 | -0.360258115 | Xaa-Pro aminopeptidase app-1; Catalyzes the removal of a penultimate prolyl residue from the N-termini of peptides, such as Arg-Pro-Pro. Has activity towards the flp-9 neuropeptide KPSFVRF- amide. (616 aa) |
| CHC-1 | P34574 | 0.042539597 | 1.371206631 | -0.368433427 | Probable clathrin heavy chain 1; Clathrin is the major protein of the polyhedral coat of coated pits and vesicles (By similarity). May play a role in yolk protein clatherin-mediated endocytosis by oocytes during oogenesis ; Belongs to the clathrin heavy chain family. (1681 aa) |
| CYN-7 | P52015 | 0.036877937 | 1.433233378 | -0.370248079 | Peptidyl-prolyl cis-trans isomerase 7; PPIases accelerate the folding of proteins. It catalyzes the cis-trans isomerization of proline imidic peptide bonds in oligopeptides; Belongs to the cyclophilin-type PPIase family. (171 aa) |
| IFA-4 | P90900 | 0.030019168 | 1.522601356 | -0.371876225 | Intermediate filament protein ifa-4; Cytoplasmic intermediate filaments provide mechanical strength to cells. Non-essential protein. (575 aa) |
| HPD-1 | Q22633 | 0.009606783 | 2.017422018 | -0.373295702 | 4-hydroxyphenylpyruvate dioxygenase; Key enzyme in the degradation of tyrosine; Belongs to the 4HPPD family. (393 aa) |
| ASP-4 | Q21966 | 0.008326527 | 2.07953613 | -0.374497927 | Aspartic protease 4; Aspartic protease, which is part of the necrosis cell death pathway. Involved in neuronal cell degeneration. Involved in heat stress response. Belongs to the peptidase A1 family. (444 aa) |
| PCP-1 | P34610 | 0.042277542 | 1.373890275 | -0.376113448 | Putative serine protease pcp-1; Belongs to the peptidase S28 family. (565 aa) |
| CGH-1 | Q95YF3 | 0.017429614 | 1.758712239 | -0.377581883 | ATP-dependent RNA helicase cgh-1; Probable RNA helicase required for oocyte and sperm function. Also required to prevent the physiological germline apoptosis mechanism killing essentially all developing oocytes. Belongs to the DEAD box helicase family. DDX6/DHH1 subfamily. |
| DKC-1 | O17919 | 0.000923838 | 3.034403964 | -0.380151851 | Predicted to enable pseudouridine synthase activity. Predicted to be involved in box H/ACA sno(s)RNA 3'-end processing and pseudouridine synthesis. Predicted to be located in nucleolus. Predicted to be part of box H/ACA snoRNP complex. Human ortholog(s) of this gene implicated in X-linked dyskeratosis congenita; aplastic anemia; and dyskeratosis congenita. Orthologous to human DKC1 (dyskerin pseudouridine synthase 1). |
| SAMS-1 | O17680 | 0.026470704 | 1.577234516 | -0.381909815 | Probable S-adenosylmethionine synthase 1; Catalyzes the formation of S-adenosylmethionine from methionine and ATP. The reaction comprises two steps that are both catalyzed by the same enzyme: formation of S-adenosylmethionine (AdoMet) and triphosphate, and subsequent hydrolysis of the triphosphate; Belongs to the AdoMet synthase family. |
| PDI-1 | Q17967 | 0.018169168 | 1.740664965 | -0.394403424 | Protein disulfide-isomerase 1; Belongs to the protein disulfide isomerase family. |
| C23G10.2 | Q10121 | 0.043012072 | 1.36640964 | -0.39470637 | RutC family protein C23G10.2. (171 aa) |
| GOT-1.2 | Q22067 | 0.032622803 | 1.486478722 | -0.397430899 | Aspartate aminotransferase, cytoplasmic; Biosynthesis of L-glutamate from L-aspartate. Important regulator of levels of glutamate, the major excitatory neurotransmitter of the central nervous system. (408 aa) |
| RAB-11.1 | O01803 | 0.022755665 | 1.64291047 | -0.410071865 | Ras-related protein rab-11.1; The small GTPases Rab are key regulators of intracellular membrane trafficking, from the formation of transport vesicles to their fusion with membranes. Rabs cycle between an inactive GDP-bound form and an active GTP-bound form that is able to recruit to membranes different set of downstream effectors directly responsible for vesicle formation, movement, tethering and fusion. Involved in regulating the meiotic maturation of oocytes. Plays a role in egg shell formation, regulating exocytosis of chondroitin proteoglycans following fertilization. |
| VHA-6 | G5EGP4 | 0.014895441 | 1.826946641 | -0.421364922 | V-type proton ATPase subunit a; Essential component of the vacuolar proton pump (V-ATPase), a multimeric enzyme that catalyzes the translocation of protons across the membranes. Required for assembly and activity of the V-ATPase. (865 aa) |
| ALDO-2 | P46563 | 0.001255996 | 2.901011633 | -0.423530582 | Fructose-bisphosphate aldolase 2. (366 aa) |
| LIN-2 | P54936 | 0.014595637 | 1.835776947 | -0.425809259 | Protein lin-2; May play a structural role in the induction of the vulva. May be required for the localization of signal transduction molecules (such as let-23 receptor) to either the basal membrane domain or the cell junctions. |
| PRDX-3 | Q21824 | 0.002602842 | 2.58455213 | -0.42611812 | Probable peroxiredoxin prdx-3; Thiol-specific peroxidase that catalyzes the reduction of hydrogen peroxide and organic hydroperoxides to water and alcohols, respectively. Plays a role in cell protection against oxidative stress by detoxifying peroxides and as sensor of hydrogen peroxide-mediated signaling events; Belongs to the peroxiredoxin family. AhpC/Prx1 subfamily. (226 aa) |
| MYO-1 | P02567 | 0.001050382 | 2.978652592 | -0.436861346 | Myosin-1; Muscle contraction. |
| Y48A6B.3 | Q9XXD4 | 0.03607244 | 1.442824478 | -0.441804009 | Putative H/ACA ribonucleoprotein complex subunit 2-like protein; Required for ribosome biogenesis. Part of a complex which catalyzes pseudouridylation of rRNA. This involves the isomerization of uridine such that the ribose is subsequently attached to C5, instead of the normal N1. Pseudouridine ('psi') residues may serve to stabilize the conformation of rRNAs (By similarity). (163 aa) |
| LBP-9 | Q965W1 | 0.046033046 | 1.336930284 | -0.445204415 | Fatty acid-binding protein homolog 9; Belongs to the calycin superfamily. Fatty-acid binding protein (FABP) family. (152 aa) |
| HSP-60 | P50140 | 0.002466453 | 2.607927167 | -0.445739546 | Chaperonin homolog Hsp-60, mitochondrial; Implicated in mitochondrial protein import and macromolecular assembly. May facilitate the correct folding of imported proteins. May also prevent misfolding and promote the refolding and proper assembly of unfolded polypeptides generated under stress conditions in the mitochondrial matrix (By similarity). (568 aa) |
| ATP-1 | Q9XXK1 | 0.006737792 | 2.171482379 | -0.44738839 | ATP synthase subunit alpha, mitochondrial; Mitochondrial membrane ATP synthase (F(1)F(0) ATP synthase or Complex V) produces ATP from ADP in the presence of a proton gradient across the membrane which is generated by electron transport complexes of the respiratory chain. F-type ATPases consist of two structural domains, F(1) - containing the extramembraneous catalytic core, and F(0) - containing the membrane proton channel, linked together by a central stalk and a peripheral stalk. |
| CYSL-2 | O45679 | 0.002076977 | 2.682568297 | -0.448366307 | Bifunctional L-3-cyanoalanine synthase/cysteine synthase; Primarily catalyzes the formation of cyanoalanine and hydrogen sulfide from cysteine and hydrogen cyanide. Can also catalyze, although less efficiently, the formation of cyanoalanine and hydrogen sulfide from either S-sulfocysteine or O-acetylserine and hydrogen cyanide and the formation of cysteine from either S-sulfocysteine or O- acetylserine and hydrogen sulfide. By catalyzing the assimilation of cyanide produced by P.aeruginosa, mediates resistance to infection. Involved in fertility, growth and aging. |
| ALH-9 | P46562 | 0.00097305 | 3.011864926 | -0.45111031 | Putative aldehyde dehydrogenase family 7 member A1 homolog; Belongs to the aldehyde dehydrogenase family. (531 aa) |
| F09E5.3 | Q19264 | 0.047928068 | 1.31941008 | -0.471042269 | Putative deoxyribose-phosphate aldolase; Catalyzes a reversible aldol reaction between acetaldehyde and D-glyceraldehyde 3-phosphate to generate 2-deoxy-D-ribose 5- phosphate. (303 aa) |
| PHB-2 | P50093 | 0.04569648 | 1.340117253 | -0.472435566 | Mitochondrial prohibitin complex protein 2; PHB proteins are essential during embryonic development and are required for somatic and germline differentiation in the larval gonad. A deficiency in PHB proteins results in altered mitochondrial biogenesis in body wall muscle cells. (294 aa) |
| COX-5A | P55954 | 0.000883667 | 3.05371125 | -0.473563871 | Cytochrome c oxidase subunit 5A, mitochondrial; Component of the cytochrome c oxidase, the last enzyme in the mitochondrial electron transport chain which drives oxidative phosphorylation. The respiratory chain contains 3 multisubunit complexes succinate dehydrogenase (complex II, CII), ubiquinol- cytochrome c oxidoreductase (cytochrome b-c1 complex, complex III, CIII) and cytochrome c oxidase (complex IV, CIV), that cooperate to transfer electrons derived from NADH and succinate to molecular oxygen, creating an electrochemical gradient over the inner membrane that drives transmembrane proton movement and ATP synthesis by ATP synthase (complex V). (174 aa) |
| RPT-4 | O17071 | 0.026877603 | 1.570609464 | -0.481591366 | Probable 26S proteasome regulatory subunit 10B; The 26S proteasome is involved in the ATP-dependent degradation of ubiquitinated proteins. The regulatory (or ATPase) complex confers ATP dependency and substrate specificity to the 26S complex (By similarity). (406 aa) |
| F27D4.1 | Q93615 | 0.022965272 | 1.638928415 | -0.481935412 | Probable electron transfer flavoprotein subunit alpha, mitochondrial; The electron transfer flavoprotein serves as a specific electron acceptor for several dehydrogenases, including five acyl-CoA dehydrogenases, glutaryl-CoA and sarcosine dehydrogenase. It transfers the electrons to the main mitochondrial respiratory chain via ETF- ubiquinone oxidoreductase (ETF dehydrogenase) (By similarity). (332 aa) |
| SODH-2 | O45687 | 0.039556248 | 1.402784911 | -0.485443413 | Alcohol dehydrogenase 2; Belongs to the zinc-containing alcohol dehydrogenase family. (351 aa) |
| LAF-1 | D0PV95;O01836 | 0.000534896 | 3.271730964 | -0.488721239 | ATP-dependent RNA helicase laf-1; Multifunctional ATP-dependent RNA helicase. Plays a role in RNA remodeling, but is not required for RNA unwinding. Binds to RNA in a concentration-dependent manner to stimulate annealing between two complementary strands of RNA. This process is also dependent upon ATP; ATP reduces binding to RNA and subsequently diminishes RNA annealing. Involved in many cellular processes, which do not necessarily require its ATPase/helicase catalytic activities. Involved in the regulation of transcription and translation initiation. Involved in innate immunity (By system. (708 aa) |
| ALDO-1 | P54216 | 0.011133189 | 1.953380424 | -0.495315374 | Fructose-bisphosphate aldolase 1. (365 aa) |
| MUA-6 | O02365 | 0.001375707 | 2.861474206 | -0.495560037 | Intermediate filament protein ifa-2; Cytoplasmic intermediate filaments provide mechanical strength to cells. Essential protein, involved in attachment structures in epidermal cells that connect muscles to the external cuticle. Probably acts by forming hypodermal hemidesmosome complexes that help mediate muscle-cuticle force transduction. Although expressed during embryogenesis, it is not required for embryonic development of muscle- cuticle linkages nor for the localization of other proteins to the hemidesmosomes in embryos. (581 aa) |
| LMTR-2 | Q9N2U6 | 0.01412265 | 1.850083812 | -0.499787826 | Ragulator complex protein LAMTOR2 homolog; Regulator of the TOR pathway, a signaling cascade that promotes cell growth in response to growth factors, energy levels, and amino acids. May activate the TOR signaling cascade in response to amino acids; Belongs to the GAMAD family. (124 aa) |
| ERFA-1 | O16520 | 0.003405471 | 2.46782286 | -0.529958315 | Eukaryotic peptide chain release factor subunit 1; Directs the termination of nascent peptide synthesis (translation) in response to the termination codons UAA, UAG and UGA. Belongs to the eukaryotic release factor 1 family. (593 aa) |
| CCT-5 | P47209 | 0.002866231 | 2.542688763 | -0.530818798 | T-complex protein 1 subunit epsilon; Molecular chaperone; assists the folding of proteins upon ATP hydrolysis. Known to play a role, in vitro, in the folding of actin and tubulin. (542 aa) |
| RPL-3 | P50880 | 0.031337567 | 1.50393472 | -0.530883529 | 60S ribosomal protein L3; The L3 protein is a component of the large subunit of cytoplasmic ribosomes; Belongs to the universal ribosomal protein uL3 family. (401 aa) |
| GPD-1 | P04970;P17331 | 0.006929133 | 2.159321092 | -0.540106884 | Glyceraldehyde-3-phosphate dehydrogenase 1. (341 aa) |
| MYO-2 | P12845 | 0.04569648 | 1.340117253 | -0.542205373 | Myosin-2; Muscle contraction. |
| OGDH-1 | O61199 | 0.009867881 | 2.005776114 | -0.543647775 | 2-oxoglutarate dehydrogenase, mitochondrial; The 2-oxoglutarate dehydrogenase complex catalyzes the overall conversion of 2-oxoglutarate to succinyl-CoA and CO(2). It contains multiple copies of three enzymatic components: 2-oxoglutarate dehydrogenase (E1), dihydrolipoamide succinyltransferase (E2) and lipoamide dehydrogenase (E3) (By similarity). (1029 aa) |
| LEC-3 | Q09581 | 0.000962531 | 3.016585155 | -0.54817155 | 32 kDa beta-galactoside-binding lectin lec-3; Binds galactose. (337 aa) |
| UCR-1 | P98080 | 0.003628914 | 2.440223329 | -0.556915496 | Cytochrome b-c1 complex subunit 1, mitochondrial. (473 aa) |
| T20H4.5 | Q22619 | 0.026026017 | 1.584592286 | -0.568268146 | Probable NADH dehydrogenase [ubiquinone] iron-sulfur protein 8, mitochondrial; Core subunit of the mitochondrial membrane respiratory chain NADH dehydrogenase (Complex I) that is believed to belong to the minimal assembly required for catalysis. Complex I functions in the transfer of electrons from NADH to the respiratory chain. The immediate electron acceptor for the enzyme is believed to be ubiquinone (By similarity). (212 aa) |
| RPL-12 | P61866 | 0.00482001 | 2.316952081 | -0.57112796 | 60S ribosomal protein L12; Binds directly to 26S ribosomal RNA; Belongs to the universal ribosomal protein uL11 family. (165 aa) |
| MAPH-1.1 | P91859 | 0.011133189 | 1.953380424 | -0.576784694 | Microtubule-associated protein homolog maph-1.1; Belongs to the MAP1A/MAP1B/MAP1S family. (878 aa) |
| CTL-2 | Q27487 | 0.010952062 | 1.9605041 | -0.577647403 | Peroxisomal catalase 1; Peroxisomal catalase involved in the oxidative stress response serving to protect cells from toxicity. Plays a role in maintaining normal lifespan. Plays a role in regulating the response to pathogens such as *E.faecalis.* (500 aa) |
| VHA-8 | Q95X44 | 0.000790486 | 3.102105648 | -0.580308738 | Vacuolar H ATPase. (226 aa) |
| C33A12.1 | Q18359 | 0.002694889 | 2.569459197 | -0.592446304 | Probable NADH dehydrogenase [ubiquinone] 1 alpha subcomplex subunit 5; Accessory subunit of the mitochondrial membrane respiratory chain NADH dehydrogenase (Complex I), that is believed not to be involved in catalysis. Complex I functions in the transfer of electrons from NADH to the respiratory chain. The immediate electron acceptor for the enzyme is believed to be ubiquinone (By similarity). (150 aa) |
| DLD-1 | O17953 | 0.014175372 | 1.848465525 | -0.593892835 | Dihydrolipoyl dehydrogenase, mitochondrial. (495 aa) |
| OSTB-1 | P45971 | 0.004709379 | 2.327036325 | -0.597201322 | Dolichyl-diphosphooligosaccharide--protein glycosyltransferase 48 kDa subunit; Subunit of the oligosaccharyl transferase (OST) complex that catalyzes the initial transfer of a defined glycan (Glc(3)Man(9)GlcNAc(2) in eukaryotes) from the lipid carrier dolichol- pyrophosphate to an asparagine residue within an Asn-X-Ser/Thr consensus motif in nascent polypeptide chains, the first step in protein N-glycosylation. (445 aa) |
| DEB-1 | P19826 | 0.030369559 | 1.517561508 | -0.602794266 | Vinculin; Involved in cell adhesion. May be involved in the attachment of the actin-based microfilaments to the plasma membrane. Involved in ovulation. (1090 aa) |
| UNC-15 | P10567 | 0.030019168 | 1.522601356 | -0.605765221 | Paramyosin; Structural component of the muscle thick filaments which is involved in assembly and organization of sarcomere myofilaments. Involved in ovulation. Plays a role in the formation of muscle connections, also called muscle arm extensions, between the body wall and the motor axons in the dorsal and ventral cord. Belongs to the paramyosin family. |
| VHA-13 | Q9XW92 | 0.010760425 | 1.968170579 | -0.612392161 | V-type proton ATPase catalytic subunit A; Catalytic subunit of the peripheral V1 complex of vacuolar ATPase (Probable). V-ATPase vacuolar ATPase is responsible for acidifying a variety of intracellular compartments in eukaryotic cells. Required along with other vacuolar ATPase components for the removal of protein aggregates which form in immature oocytes in the distal gonad. This removal occurs as the oocytes mature and move to the proximal gonad, is triggered by the introduction of sperm through mating and occurs before fertilization. (606 aa) |
| CYN-2 | P52010 | 0.001050382 | 2.978652592 | -0.616221178 | Peptidyl-prolyl cis-trans isomerase 2; PPIases accelerate the folding of proteins. It catalyzes the cis-trans isomerization of proline imidic peptide bonds in oligopeptides; Belongs to the cyclophilin-type PPIase family. (172 aa) |
| IFD-1 | Q86DC6 | 0.013020324 | 1.885378225 | -0.622329887 | Intermediate filament protein ifd-1; Cytoplasmic intermediate filaments provide mechanical strength to cells. Not essential protein. (575 aa) |
| PBS-3 | Q23237 | 0.00973062 | 2.011859494 | -0.622715376 | Proteasome subunit beta type-3; The proteasome is a multicatalytic proteinase complex which is characterized by its ability to cleave peptides with Arg, Phe, Tyr, Leu, and Glu adjacent to the leaving group at neutral or slightly basic pH. The proteasome has an ATP-dependent proteolytic activity (By similarity). (204 aa) |
| SDHA-1 | Q09508 | 0.027853675 | 1.555117495 | -0.623440714 | Succinate dehydrogenase [ubiquinone] flavoprotein subunit, mitochondrial; Flavoprotein (FP) subunit of succinate dehydrogenase (SDH) that is involved in complex II of the mitochondrial electron transport chain and is responsible for transferring electrons from succinate to ubiquinone (coenzyme Q); Belongs to the FAD-dependent oxidoreductase 2 family. FRD/SDH subfamily. (646 aa) |
| RPS-12 | P49196 | 0.019526525 | 1.709375044 | -0.626353283 | 40S ribosomal protein S12; Belongs to the eukaryotic ribosomal protein eS12 family. (140 aa) |
| GFM-1 | Q9XV52 | 0.034109125 | 1.467129419 | -0.633610372 | Elongation factor G, mitochondrial; Mitochondrial GTPase that catalyzes the GTP-dependent ribosomal translocation step during translation elongation. During this step, the ribosome changes from the pre-translocational (PRE) to the post-translocational (POST) state as the newly formed A-site-bound peptidyl-tRNA and P-site-bound deacylated tRNA move to the P and E sites, respectively. Catalyzes the coordinated movement of the two tRNA molecules, the mRNA and conformational changes in the ribosome. (750 aa) |
| GPD-2 | P17329;P17330 | 0.002997066 | 2.523303624 | -0.636822882 | Glyceraldehyde-3-phosphate dehydrogenase 2. (341 aa) |
| LEC-1 | P36573 | 0.02539624 | 1.595230584 | -0.641906662 | 32 kDa beta-galactoside-binding lectin; Binds galactose. (285 aa) |
| UNC-52 | Q06561 | 0.006929133 | 2.159321092 | -0.647271179 | Basement membrane proteoglycan; Component of an integrin containing attachment complex, which is required for muscle development and maintenance. Probable structural role in myofilament assembly and/or attachment of the myofilament lattice to the cell membrane. May be an extracellular anchor for integrin receptors in body wall muscles and myoepithelial sheath cells. During the formation of neuromuscular junctions at the larval stage, negatively regulates membrane protrusion from body wall muscles, probably downstream of the integrin complex formed by pat-2 and pat-3. Involved in ovulation. |
| RPL-21 | P34334 | 0.042547612 | 1.37112481 | -0.647554577 | 60S ribosomal protein L21; Belongs to the eukaryotic ribosomal protein eL21 family. (161 aa) |
| ASP-1 | G5EEI4 | 0.001450753 | 2.838406536 | -0.647790073 | Aspartic protease 1; Aspartic protease, which is part of the necrosis cell death pathway. Promotes B.thuringiensis Cry6Aa stability by preventing its proteolysis by host gut proteases. Required for Cry6Aa-induced necrotic death of intestinal cells. Cry6Aa uptake into the host intestinal cells triggers an increase in intracellular Ca(2+) levels leading to lysosome rupture and to the subsequent release of asp-1 which leads to necrosis ; Belongs to the peptidase A1 family. |
| ALH-8 | P52713 | 0.000420359 | 3.376379237 | -0.663024794 | Probable methylmalonate-semialdehyde dehydrogenase [acylating], mitochondrial; Plays a role in valine and pyrimidine metabolism. Binds fatty acyl-CoA (By similarity); Belongs to the aldehyde dehydrogenase family. (523 aa) |
| VDAC-1 | Q21752 | 0.014175372 | 1.848465525 | -0.664515755 | Probable voltage-dependent anion-selective channel; Forms a channel through the cell membrane that allows diffusion of small hydrophilic molecules. (283 aa) |
| GSNL-1 | Q21253 | 0.011133189 | 1.953380424 | -0.667382358 | Gelsolin-like protein 1; Calcium-regulated, actin-modulating protein that binds to the plus (or barbed) ends of actin monomers or filaments, preventing monomer exchange (end-blocking or capping). Binds actin but does not nucleate actin polymerization, albeit slows down elongation by blocking the barbed ends. By promoting actin depolymerization, required for the elimination of presynaptic components downstream of the egl-1, ced-4 and ced-3 apoptotic pathway during larval development. Belongs to the villin/gelsolin family. (475 aa) |
| SUCA-1 | P53588 | 0.016600095 | 1.779889424 | -0.66847367 | Succinate--CoA ligase [ADP-forming] subunit beta, mitochondrial; ATP-specific succinyl-CoA synthetase functions in the citric acid cycle (TCA), coupling the hydrolysis of succinyl-CoA to the synthesis of ATP and thus represents the only step of substrate-level phosphorylation in the TCA. The beta subunit provides nucleotide specificity of the enzyme and binds the substrate succinate, while the binding sites for coenzyme A and phosphate are found in the alpha subunit. (435 aa) |
| DNPP-1 | Q19087 | 0.001850123 | 2.732799292 | -0.685169378 | Aspartyl aminopeptidase; Aminopeptidase with specificity towards an acidic amino acid at the N-terminus. Plays a role in membrane trafficking and is specifically involved in the recycling and degradation of endocytic cargo. (507 aa) |
| PRP-8 | P34369 | 0.030992678 | 1.5087409 | -0.698237388 | Pre-mRNA-splicing factor 8 homolog; Functions as a scaffold that mediates the ordered assembly of spliceosomal proteins and snRNAs. Required for the assembly of the U4/U6-U5 tri-snRNP complex. Functions as scaffold that positions spliceosomal U2, U5 and U6 snRNAs at splice sites on pre-mRNA substrates, so that splicing can occur. Interacts with both the 5' and the 3' splice site. (2329 aa) |
| CCT-3 | Q9N4J8 | 0.046519271 | 1.332367102 | -0.699735826 | T-complex protein 1 subunit gamma; Molecular chaperone; assists the folding of proteins upon ATP hydrolysis (By similarity). Known to play a role, in vitro, in the folding of actin and tubulin (By similarity). Plays a role in microtubule polymerization. (543 aa) |
| CYN-3 | P52011 | 0.039367421 | 1.404863036 | -0.708819676 | Peptidyl-prolyl cis-trans isomerase 3; PPIases accelerate the folding of proteins. It catalyzes the cis-trans isomerization of proline imidic peptide bonds in oligopeptides. (173 aa) |
| DIM-1 | Q18066 | 0.001305572 | 2.884199286 | -0.711258951 | Disorganized muscle protein 1; Stabilizes the attachment of the myofilament lattice to the muscle cell membrane. Isoform b is sufficient for wild-type function. (640 aa) |
| ANC-1 | Q9N4M4 | 0.011263892 | 1.948311523 | -0.721681792 | Nuclear anchorage protein 1; Plays a central role in nuclear and mitochondrial anchoring. Probably connects nuclei to the cytoskeleton by interacting with unc-84 at the nuclear envelope and with F-actin in the cytoplasm, creating a bridge across the nuclear envelope between the cytoskeleton and the nucleus. Has a role in positioning of the cell body of the PVQ lumbar interneuron. Belongs to the nesprin family. |
| RPL-9 | Q95Y90 | 0.000815301 | 3.088681998 | -0.725128519 | 60S ribosomal protein L9; Belongs to the universal ribosomal protein uL6 family. (189 aa) |
| ELO-1 | G5EEE5 | 0.002694889 | 2.569459197 | -0.734344928 | Elongation of very long chain fatty acids protein. (288 aa) |
| RPL-11.1 | Q94300 | 0.008894539 | 2.050876535 | -0.744800157 | 60S ribosomal protein L11-1; Component of the ribosome, a large ribonucleoprotein complex responsible for the synthesis of proteins in the cell. The small ribosomal subunit (SSU) binds messenger RNAs (mRNAs) and translates the encoded message by selecting cognate aminoacyl-transfer RNA (tRNA) molecules. The large subunit (LSU) contains the ribosomal catalytic site termed the peptidyl transferase center (PTC), which catalyzes the formation of peptide bonds, thereby polymerizing the amino acids delivered by tRNAs into a polypeptide chain. (196 aa) |
| CTC-2 | P24894 | 0.009352795 | 2.029058572 | -0.772003023 | Cytochrome c oxidase subunit 2; Component of the cytochrome c oxidase, the last enzyme in the mitochondrial electron transport chain which drives oxidative phosphorylation. The respiratory chain contains 3 multisubunit complexes succinate dehydrogenase (complex II, CII), ubiquinol- cytochrome c oxidoreductase (cytochrome b-c1 complex, complex III, CIII) and cytochrome c oxidase (complex IV, CIV), that cooperate to transfer electrons derived from NADH and succinate to molecular oxygen, creating an electrochemical gradient over the inner membrane that drives transmembrane transport. (231 aa) |
| MCCC-2 | P34385 | 0.036396041 | 1.438945855 | -0.777094203 | Carboxyltransferase subunit of the 3-methylcrotonyl-CoA carboxylase, an enzyme that catalyzes the conversion of 3-methylcrotonyl-CoA to 3-methylglutaconyl-CoA, a critical step for leucine and isovaleric acid catabolism. |
| DPYD-1 | Q18164 | 0.021748596 | 1.662568781 | -0.781168998 | Dihydropyrimidine dehydrogenase [NADP(+)]; Involved in pyrimidine base degradation. Catalyzes the reduction of uracil and thymine (By similarity). Involved in the degradation of the chemotherapeutic drug 5-fluorouracil. (1059 aa) |
| CTSA-3.1 | P52716 | 0.000636875 | 3.195945591 | -0.781478059 | Serine carboxypeptidase ctsa-3.1; Belongs to the peptidase S10 family. (574 aa) |
| PYR-1 | Q18990 | 0.001157779 | 2.936374518 | -0.783008257 | Glutamine-dependent carbamoyl-phosphate synthase; This protein is a 'fusion' protein encoding four enzymatic activities of the pyrimidine pathway (GATase, CPSase, ATCase and DHOase) (By similarity). Involved in the elongation of the pharyngeal isthmus during development, probably by providing precursors of UDP- sugars required for heparan sulfate proteoglycan biosynthesis. Regulates the organization of the actin and intermediate filaments cytoskeleton in the pharyngeal muscles ; In the central section; belongs to the metallo-dependent hydrolases superfamily. DHOase family. CAD subfamily. (2198 aa) |
| VHA-5 | G5EEK9 | 0.00097305 | 3.011864926 | -0.787970028 | V-type proton ATPase subunit a; Essential component of the vacuolar proton pump (V-ATPase), a multimeric enzyme that catalyzes the translocation of protons across the membranes. Required for assembly and activity of the V-ATPase. (873 aa) |
| MMCM-1 | Q23381 | 0.018193242 | 1.740089909 | -0.793283842 | Probable methylmalonyl-CoA mutase, mitochondrial; Involved, in man, in the degradation of several amino acids, odd-chain fatty acids and cholesterol via propionyl-CoA to the tricarboxylic acid cycle. MCM has different functions in other species (By similarity). (744 aa) |
| IFC-2 | Q21067 | 0.015129753 | 1.82016816 | -0.793846522 | Intermediate filament protein ifc-2; Cytoplasmic intermediate filaments provide mechanical strength to cells. Not essential protein, although its absence leads to mild defects in locomotion. (1248 aa) |
| LAM-2 | Q18823 | 0.009482245 | 2.023088823 | -0.79526496 | Laminin-like protein lam-2; During the formation of neuromuscular junctions at the larval stage, negatively regulates membrane protrusion from body wall muscles, probably downstream of the integrin complex formed by pat-2 and pat-3. |
| PAS-5 | Q95008 | 0.000884933 | 3.053089463 | -0.808387627 | Proteasome subunit alpha type-5; The proteasome is a multicatalytic proteinase complex which is characterized by its ability to cleave peptides with Arg, Phe, Tyr, Leu, and Glu adjacent to the leaving group at neutral or slightly basic pH. The proteasome has an ATP-dependent proteolytic activity (By similarity). (248 aa) |
| FIB-1 | Q22053 | 0.007037511 | 2.152580899 | -0.820201465 | rRNA 2'-O-methyltransferase fibrillarin; S-adenosyl-L-methionine-dependent methyltransferase that has the ability to methylate both RNAs and proteins (By similarity). Involved in pre-rRNA processing. Utilizes the methyl donor S-adenosyl- L-methionine to catalyze the site-specific 2'-hydroxyl methylation of ribose moieties in pre-ribosomal RNA (By similarity). Site specificity is provided by a guide RNA that base pairs with the substrate. Methylation occurs at a characteristic distance from the sequence involved in base pairing with the guide RNA (By similarity). |
| RPL-11.2 | Q19162 | 0.001157779 | 2.936374518 | -0.831466295 | 60S ribosomal protein L11-2; Component of the ribosome, a large ribonucleoprotein complex responsible for the synthesis of proteins in the cell. The small ribosomal subunit (SSU) binds messenger RNAs (mRNAs) and translates the encoded message by selecting cognate aminoacyl-transfer RNA (tRNA) molecules. The large subunit (LSU) contains the ribosomal catalytic site termed the peptidyl transferase center (PTC), which catalyzes the formation of peptide bonds, thereby polymerizing the amino acids delivered by tRNAs into a polypeptide chain. (196 aa) |
| RACK-1 | Q21215 | 0.019103772 | 1.718880879 | -0.834169912 | Guanine nucleotide-binding protein subunit beta-2-like 1; Required for the expression of antimicrobial peptide nlp-29 in response to fungal infection or physical injury. (325 aa) |
| ACOX-1.1 | O62140;O62138 | 0.002866231 | 2.542688763 | -0.868646116 | Acyl-coenzyme A oxidase; Belongs to the acyl-CoA oxidase family. (674 aa) |
| IDHA-1 | Q93714 | 0.027923968 | 1.554022871 | -0.86936075 | Probable isocitrate dehydrogenase [NAD] subunit alpha, mitochondrial; Belongs to the isocitrate and isopropylmalate dehydrogenases family. (358 aa) |
| LDP-1 | Q9GZE9 | 0.002113853 | 2.674925226 | -0.880528417 | Lipid droplet localized protein; Belongs to the saccharopine dehydrogenase family. (426 aa) |
| GLH-4 | O76743 | 0.004373503 | 2.359170574 | -0.881574087 | ATP-dependent RNA helicase glh-4; Probable ATP-binding RNA helicase. May act redundantly with the P-granule component glh-1 to regulate the formation of the granular structure of P-granules in embryos. May protect somatic cells from excessive apoptosis during normal development. |
| PAS-1 | O17586 | 0.015504571 | 1.809540236 | -0.882376734 | Proteasome subunit alpha type-6; The proteasome is a multicatalytic proteinase complex which is characterized by its ability to cleave peptides with Arg, Phe, Tyr, Leu, and Glu adjacent to the leaving group at neutral or slightly basic pH. The proteasome has an ATP-dependent proteolytic activity (By similarity). |
| NKB-3 | Q9XUY5 | 0.001500045 | 2.823895782 | -0.914442519 | Probable sodium/potassium-transporting ATPase subunit beta-3; This is the non-catalytic component of the active enzyme, which catalyzes the hydrolysis of ATP coupled with the exchange of Na(+) and K(+) ions across the plasma membrane. The beta subunit regulates, through assembly of alpha/beta heterodimers, the number of sodium pumps transported to the plasma membrane (By similarity). Implicated in genomic response to various soil bacteria that affects fitness, lifespan and brood size. (317 aa) |
| MDH-1 | Q9UAV5 | 6.06261E-05 | 4.21734063 | -0.925683562 | Malate dehydrogenase. (336 aa) |
| ECH-6 | P34559 | 0.012583585 | 1.900195612 | -0.927628989 | Probable enoyl-CoA hydratase, mitochondrial. (288 aa) |
| TTR-16 | P55955 | 0.039367421 | 1.404863036 | -0.949908431 | Transthyretin-like protein 16. (132 aa) |
| EMB-9 | P17139 | 0.029989705 | 1.523027809 | -0.969814161 | Collagen alpha-1(IV) chain; Collagen type IV is specific for basement membranes (Probable). Required to restrict presynaptic growth at the neuromuscular junctions (NMJ) in late larval stage and in adult motor neurons. May play a role in axon regeneration in embryos following injury in D-type motor neurons. |
| RPL-13A | Q27389 | 0.009984152 | 2.000688816 | -0.973600669 | rpl-13A encodes large ribosomal subunit protein uL13, a protein of the large ribosomal subunit. |
| EIF-2BETA | Q21230 | 0.018169168 | 1.740664965 | -0.984649825 | Eukaryotic translation initiation factor 2 subunit 2; eIF-2 functions in the early steps of protein synthesis by forming a ternary complex with GTP and initiator tRNA. This complex binds to a 40S ribosomal subunit, followed by mRNA binding to form a 43S preinitiation complex. Junction of the 60S ribosomal subunit to form the 80S initiation complex is preceded by hydrolysis of the GTP bound to eIF-2 and release of an eIF-2-GDP binary complex. (250 aa) |
| PCCA-1 | Q19842 | 0.001585071 | 2.799951169 | -1.006020612 | Propionyl-CoA carboxylase alpha chain, mitochondrial; This is one of the 2 subunits of the biotin-dependent propionyl-CoA carboxylase (PCC), a mitochondrial enzyme involved in the catabolism of odd chain fatty acids, branched-chain amino acids isoleucine, threonine, methionine, and valine and other metabolites. Propionyl-CoA carboxylase catalyzes the carboxylation of propionyl- CoA/propanoyl-CoA to D-methylmalonyl-CoA/(S)-methylmalonyl-CoA (By similarity). (724 aa) |
| AC3.5 | Q17405 | 0.001585071 | 2.799951169 | -1.01118205 | Aminopeptidase-like protein AC3.5. (1090 aa) |
| RPS-6 | Q9NEN6 | 0.000720613 | 3.142297676 | -1.095119899 | 40S ribosomal protein S6; May play an important role in controlling cell growth and proliferation through the selective translation of particular classes of mRNA (By similarity). Negatively regulates lifespan ; Belongs to the eukaryotic ribosomal protein eS6 family. (246 aa) |
| RPS-0 | P46769 | 0.000312402 | 3.505285981 | -1.104078126 | 40S ribosomal protein SA; Required for the assembly and/or stability of the 40S ribosomal subunit. Required for the processing of the 20S rRNA- precursor to mature 18S rRNA in a late step of the maturation of 40S ribosomal subunits. (276 aa) |
| CYN-1 | P52009 | 0.001305572 | 2.884199286 | -1.127043275 | Peptidyl-prolyl cis-trans isomerase 1; PPIases accelerate the folding of proteins. It catalyzes the cis-trans isomerization of proline imidic peptide bonds in oligopeptides; Belongs to the cyclophilin-type PPIase family. (192 aa) |
| B0272.3 | P41938 | 0.029662907 | 1.527786285 | -1.128909413 | Probable 3-hydroxyacyl-CoA dehydrogenase B0272.3; Belongs to the 3-hydroxyacyl-CoA dehydrogenase family. (309 aa) |
| KAT-1 | Q22100 | 0.013278698 | 1.87684451 | -1.129564853 | Acetyl-CoA acetyltransferase homolog, mitochondrial; Plays a role in ketone body metabolism (By similarity). Has an effect on lysosome related organelles (LRO) function, in a pathway with serotonin. Acts downstream of sir-2.1 to regulate aging and stress resistance. Belongs to the thiolase-like superfamily. Thiolase family. (407 aa) |
| PDI-6 | Q11067 | 0.013020324 | 1.885378225 | -1.140236419 | Protein disulfide-isomerase A6 homolog; May function as a chaperone that inhibits aggregation of misfolded proteins (By similarity). May negatively regulate the unfolded protein response (UPR) through binding to UPR sensors ; Belongs to the protein disulfide isomerase family. (440 aa) |
| CPL-1 | O45734 | 0.006140128 | 2.211822579 | -1.141719372 | Cathepsin L-like; Cysteine protease which plays an essential role in the degradation of proteins in lysosomes. During early embryogenesis, maternally required for the proteolytic processing of yolk proteins in platelets, a lysosome- like structure where a slow and controlled degradation of yolk proteins occurs. In the gonad, required for the clearance of apoptotic germ cells in the engulfing cell phagolysosomes. In embryos, required for the degradation of endocytic and autophagic cargos. In embryos, may play a role in the degradation of lipid-containing droplets. |
| CUL-4 | Q17392 | 0.028409888 | 1.546530478 | -1.177877525 | Cullin-4; Component of cullin-based E3 ubiquitin-protein ligase complexes which mediate the ubiquitination and subsequent proteasomal degradation of target proteins. The functional specificity of the E3 ubiquitin-protein ligase complex depends on the variable substrate recognition component. In association with ddb-1 directs ubiquitination of cdt-1 during S phase and is required for restraining DNA rereplication. Probably is involved in ubiquitination of cki-1. (840 aa) |
| EPI-1 | Q21313 | 0.034198595 | 1.465991736 | -1.179064804 | Laminin-like protein epi-1; During the formation of neuromuscular junctions at the larval stage, negatively regulates membrane protrusion from body wall muscles, probably downstream of the integrin complex formed by pat-2 and pat-3. |
| LYS-1 | O62415 | 0.000317787 | 3.497863998 | -1.179927889 | Lysozyme-like protein 1; Involved in resistance to Gram-negative bacterium S.marcescens and to bacterium Gram-positive S.aureus infection. |
| MSP-19 | P53017;P05634;P53018;Q18461;Q21244;Q23519;P53019;P53020;Q9TVW5;Q17856;Q19832;Q94053 | 0.001103621 | 2.957179902 | -1.183283824 | Major sperm protein 19/31/40/45/50/51/53/59/61/65/81/113/142. |
| TOMM-40 | Q18090 | 0.006109305 | 2.214008214 | -1.184826411 | Mitochondrial import receptor subunit TOM40 homolog; Channel-forming protein essential for import of protein precursors into mitochondria. Specifically required for nnt-1 accumulation in the mitochondria and may be involved in the secretion of daf-28/insulin from the mitochondria. Required for embryonic and larval development. (301 aa) |
| DARS-1 | Q03577 | 0.00040242 | 3.395320421 | -1.287698635 | Aspartate--tRNA ligase, cytoplasmic. (531 aa) |
| PAS-2 | Q27488 | 0.002707351 | 2.567455395 | -1.304374872 | Proteasome subunit alpha type-2; The proteasome is a multicatalytic proteinase complex which is characterized by its ability to cleave peptides with Arg, Phe, Tyr, Leu, and Glu adjacent to the leaving group at neutral or slightly basic pH. The proteasome has an ATP-dependent proteolytic activity (By similarity). (231 aa) |
| MYO-3 | P12844 | 0.000802338 | 3.095642854 | -1.376715999 | Myosin-3; Essential for muscle contraction. Involved in ovulation likely by regulating the contraction of gonadal myoepithelial sheath cells. Belongs to the TRAFAC class myosin-kinesin ATPase superfamily. Myosin family. |
| TARS-1 | P52709 | 0.002694889 | 2.569459197 | -1.378551562 | Threonine--tRNA ligase, cytoplasmic; Belongs to the class-II aminoacyl-tRNA synthetase family. (725 aa) |
| DBT-1 | Q23571 | 0.004373503 | 2.359170574 | -1.451276183 | 2-oxoisovalerate dehydrogenase E2 component (dihydrolipoyl transacylase); The branched-chain alpha-keto dehydrogenase complex catalyzes the overall conversion of alpha-keto acids to acyl-CoA and CO(2) (By similarity). It contains multiple copies of three enzymatic components: branched-chain alpha-keto acid decarboxylase (E1), lipoamide acyltransferase (E2) and lipoamide dehydrogenase (E3) (By similarity). (448 aa) |
| NSUN-1 | Q9TYV5 | 0.00097305 | 3.011864926 | -1.462925979 | NSUN-1 is an RNA methyltransferase belonging to the NOP2/Sun (NSUN) family. It catalyzes the methylation of cytosine residues in RNA, specifically the formation of 5-methylcytosine (m⁵C). In *C. elegans*, NSUN-1 modifies 26S rRNA at position C2982. |
| ASP-6 | O01530 | 0.000317787 | 3.497863998 | -1.586880954 | Aspartic protease 6; Aspartic protease; Belongs to the peptidase A1 family. (389 aa) |
| VHA-19 | Q9TYW1 | 0.000534896 | 3.271730964 | -1.683482708 | Vacuolar H ATPase. (451 aa) |
| RPN-11 | O76577 | 0.033739024 | 1.471867481 | -2.229337176 | 26S proteasome non-ATPase regulatory subunit 14; Metalloprotease component of the 26S proteasome that specifically cleaves 'Lys-63'-linked polyubiquitin chains. The 26S proteasome is involved in the ATP-dependent degradation of ubiquitinated proteins. The function of the 'Lys-63'-specific deubiquitination of the proteasome is unclear (By similarity). (312 aa) |
| NFI-1 | Q5H9N3 | 0.008326527 | 2.07953613 | -2.284721285 | CTF/NF-I domain-containing protein. (823 aa) |
| MLK-1 | A0A0K3AV08 | 0.00651021 | 2.186404993 | -2.889847486 | Mitogen-activated protein kinase kinase kinase mlk-1; Serine/threonine-protein kinase which, by phosphorylating and activating mek-1, plays an important role in the activation of the JNK pathway composed of mlk-1, mek-1 and kgb-1. Involved in the response to environmental stress such as heavy metals. By activating the JNK pathway downstream of tyrosine receptor svh-2, plays a role in axon regeneration after injury. |

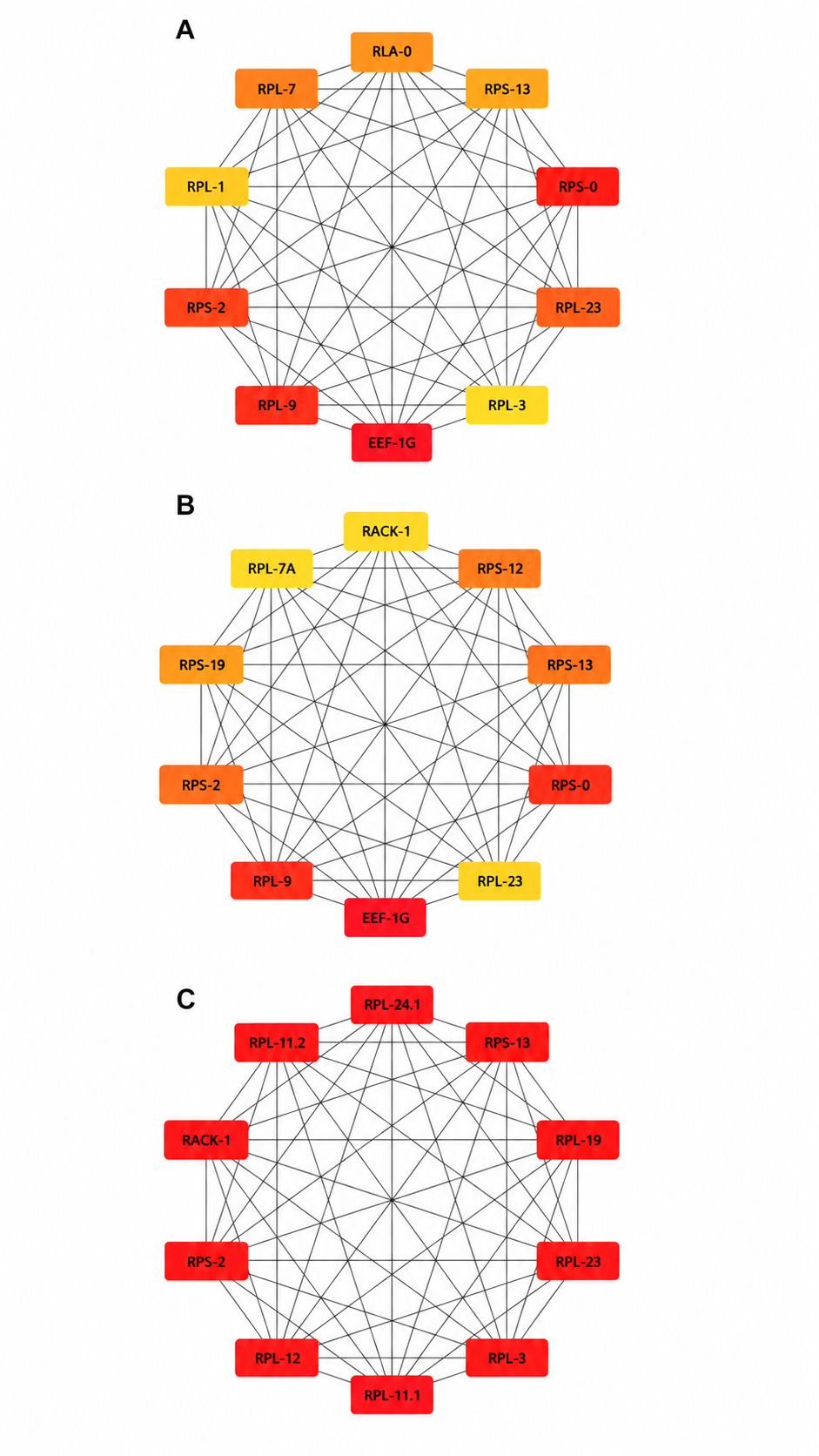

**Figure S2. Identification of hub proteins in the protein–protein interaction (PPI) network of differentially expressed proteins in *C. elegans* following CBZ exposure.** The top 10 hub proteins were identified using the CytoHubba plugin in Cytoscape based on three topological algorithms: (A) Closeness Centrality (Closeness), (B) Degree Centrality (Degree), and (C) Maximal Clique Centrality (MCC). Nodes represent proteins and edges represent predicted protein–protein interactions. Node color intensity reflects the ranking score within each algorithm, with red indicating higher-ranked hub proteins and yellow indicating lower-ranked proteins. The three algorithms consistently identified ribosomal proteins and the translation elongation factor EEF-1G as the principal hubs, highlighting the protein synthesis machinery as the most interconnected functional module affected by 0.01 mg.mL^-1^ CBZ exposure.
